# Organ-specific rewiring of mitochondrial integrity through COX7A dictates cellular ploidy control

**DOI:** 10.1101/2025.09.29.678879

**Authors:** Archan Chakraborty, Sophia DeLuca, Meera Gangasani, Stephen Rogers, Nenad Bursac, Donald T. Fox

## Abstract

To achieve proper cell and tissue size, cytoplasmic and nuclear growth must be coordinated. Disrupting this coordination causes birth defects and disease. In nature’s largest cells, nuclear growth occurs through polyploidization (whole-genome-duplication). How the massive nuclear growth of polyploid cells is coordinated with cytoplasmic growth processes such as mitochondrial biogenesis is relatively unclear. Here, focusing on one of nature’s most commonly polyploid organs-the heart-we uncover cross-talk between cytoplasmic mitochondrial biogenesis/integrity and nuclear growth/polyploidy. From a human-to-fly screen, we uncover novel regulators of cardiomyocyte ploidy, including mitochondrial integrity regulators. In comparing these cardiac hits with a parallel screen in another polyploid tissue, the salivary gland, we discovered two opposing roles for Cytochrome-c-oxidase-subunit-7A (COX7A). While salivary gland COX7A preserves mitochondrial integrity to promote polyploidy and optimal organ growth, cardiac COX7A instead suppresses mitochondrial biogenesis to repress polyploidy and prevent hypertrophic organ growth. Among all electron transport chain genes, only COX7A functions as a cardiac growth repressor. Fly hearts with compromised COX7A show abnormally high cardiac output. Human COX7A1, a mitochondrial-localized protein, similarly represses polyploidy of human iPSC-derived cardiomyocytes. In summary, our human-fly-human approach reveals conserved rewiring of mitochondrial integrity in heart tissue that switches COX7A’s role from ploidy promotion to repression. Our findings reveal fundamental cross-talk between mitochondrial biogenesis and genome duplication that are critical in growing metazoan tissues.

## INTRODUCTION

Control of cellular size during organ growth underlies all organ formation events. Two important aspects of cell size control are: 1) coordination of nuclear and cytoplasm growth (Hayden et al., 2022; Jukam et al., 2021; Li et al., 2023; Pierconti et al., 2018; Shindo and Amodeo, 2021), and 2) control of optimal cell size. In addition to coordinating the amount of cytoplasmic organelle material with genome content in the nucleus, optimal cell size in each organ must also be carefully regulated. For example, the structure and/or function of many organs can be disrupted in conditions of cellular enlargement (hypertrophy) (Blasi and Klotman, 2023; Frade and López-Sánchez, 2017; Kim et al., 2022; Sukhacheva et al., 2025). As such, there is a pressing need to understand mechanisms that achieve proper organ growth and function through regulation of optimal cellular size.

A major mechanism that drives organ growth during development is whole genome duplication, also known as polyploidization (Darmasaputra et al., 2024; Morris et al., 2024; Orr-Weaver, 2015; Pennisi, 2023). Polyploidization frequently involves endoreplication, whereby cells replicate their DNA without undergoing complete cell division (Almeida Machado Costa et al., 2022; Frawley and Orr-Weaver, 2015). This substantial increase in genome content (ploidy) massively increases nuclear size (Heijo et al., 2020) and thus cell size, which achieves higher overall organ growth (Cadart and Heald, 2022; Morris et al., 2024; Yahya et al., 2022). While polyploidization is conserved across a wide range of metazoan organs, including the heart, liver, placenta, and others (Carriere, 1969; Ebert and Pfitzer, 1977; Velicky et al., 2018; Winkelmann et al., 1987), it is especially common in the myocytes of the heart (cardiomyocytes) (Chakraborty et al., 2023; Derks and Bergmann, 2020; Han et al., 2020; Patterson et al., 2017; Soonpaa et al., 1996).

Cardiomyocytes exhibit tightly regulated ploidy levels, with both hypo-and hyper-polyploidization linked to altered cell and organ size and impaired heart function (Chakraborty et al., 2023; Elia et al., 2023). The cellular mechanisms governing the developmental transition to polyploidy are beginning to be characterized in cardiomyocytes (Derks and Bergmann, 2020; Edgar et al., 2014; Gan et al., 2020; Kyösola et al., 1988; Lan et al., 2025). However, the molecular pathways that coordinate the extent of nuclear material increase (ploidy) and cytoplasmic material increase (e.g. mitochondrial biogenesis) to achieve optimal myocardium structure still remain largely unclear.

In the cytoplasm, mitochondrial regulation appears to be linked to polyploidy regulation (Ma et al., 2011; Roh et al., 2012). Mitochondrial ATP production is essential for driving endoreplication (Ma et al., 2011). However, an increase in the ADP:ATP ratio is associated with cardiac hypertrophy, where cardiomyocytes exhibit unscheduled polyploidization (Bischof et al., 2021). Further, cardiomyocytes exhibit unique mitochondrial wiring. For example, the heart and brain have differences in the assembly of mitochondrial complex I, and these differences are thought to underly cardiac conditions such as cardiac hypertrophy (Nicol et al., 2023).

*Drosophila* provides an accessible in vivo model to discover conserved principles of cardiac development (Martínez-Morentin et al., 2015; Wessells et al., 2004; Wolf et al., 2006; Yu et al., 2013; Yu et al., 2015; Zechini et al., 2022). In our previous work, we demonstrated that cardiomyocytes in the *Drosophila* larval heart exhibit higher ploidy and cell size than those in the atrial aorta, which drives formation of a larger cardiac chamber. We observed a similar ploidy asymmetry between the left atria and ventricle of human organ donor hearts (Chakraborty et al., 2023). In both flies and humans, we found the higher ploidy chamber exhibits higher levels of insulin signaling, which is required for cardiac chamber ploidy differences in flies. This chamber-specific asymmetry in cardiomyocyte polyploidy plays a vital role in maintaining cardiac function in *Drosophila*, suggesting that different regions within the same organ may undergo specific modulation of genome content and ultimately organ growth to support structural or functional demands.

Building on our previous findings, here we conducted a targeted human-to-fly reverse genetic screen in the *Drosophila* larval heart, focusing on genes that are highly expressed in polyploid human ventricular cardiomyocytes compared to their lower-ploidy atrial counterparts. This approach led us to identify multiple novel cardiomyocyte ploidy and organ growth regulators. Interestingly, many genes identified in our cardiac screen do not similarly influence ploidy in another polyploid tissue, the salivary gland. Most strikingly, *Cytochrome c oxidase subunit 7A* (*COX7A*) represses ploidy and organ growth in the heart but promotes these same processes in the salivary gland. COX7A orthologs function in mitochondrial biology (Feng et al., 2022; García-Poyatos et al., 2024). Here, we find that cardiac COX7A does not influence mitochondrial integrity but instead suppresses mitochondrial biogenesis. This mitochondrial repression blocks positive feedback between cytoplasmic growth (mitochondrial biogenesis) and nuclear growth (polyploidy), enabling the heart to achieve optimal cell and organ size. In contrast, in the salivary gland, COX7A promotes mitochondrial integrity and biogenesis, which drives optimal cell and organ size by fueling both cellular (mitochondrial) and nuclear (ploidy) growth. COX7A is the only annotated electron transport chain regulator that functions as a cardiac growth repressor. Building on our findings in flies, we show that human mitochondrial COX7A1 also suppresses polyploidy in human induced pluripotent stem cell-derived cardiomyocytes (hiPSC-CMs). Overall, our findings reveal an organ-specific rewiring of mitochondrial control through COX7A that dictates ploidy-based growth control. Our work is significant given that dysregulated cardiomyocyte ploidy is frequently associated with pathological conditions in the human heart, including heart failure and cardiomyopathies (Beltrami et al., 1997; Derks and Bergmann, 2020; Gan et al., 2020; Swynghedauw and Delcayre, 1982).

## RESULTS

### A human-to-fly screen identifies novel cardiomyocyte ploidy regulators

Given our prior work on chamber-specific differences in cardiac ploidy, we reasoned that genes that are differentially expressed between chambers may represent ploidy regulators. Indeed, our prior analysis of single nuclear RNA-seq differences between human ventricular (vs. atrial) cardiomyocytes led to the finding that insulin receptor levels control cardiac chamber-specific polyploidy in *Drosophila* (Chakraborty et al., 2023; Litvinukova et al., 2020; Russell et al., 2025). This finding demonstrated that *Drosophila* can be used to mine human chamber-specific expression data to reveal new cardiac developmental phenotypes. Building on our findings with insulin signaling, we performed a reverse genetic screen of 92 *Drosophila* orthologs of genes that are enriched in human ventricles versus atria (**Fig 1A; TableS1: “Tab1-Ploidy Screen”)**. We used the cardiomyocyte driver *NP5164-Gal4* (hereafter: *NP>*) (Chakraborty et al., 2023), to express UAS-RNA interference (RNAi) constructs in the developing larval *Drosophila* heart (**Fig 1A**). We previously established that endoreplication and NP> expression initiate in the first larval instar after hatching (**Fig 1B**), thereby avoiding inducing phenotypes prior to cardiac endoreplication. We thus assayed final cardiomyocyte ploidy in control and RNAi animals in wandering 3^rd^ instar larvae (WL3), after the completion of the endoreplication phase.

**Figure 1:**
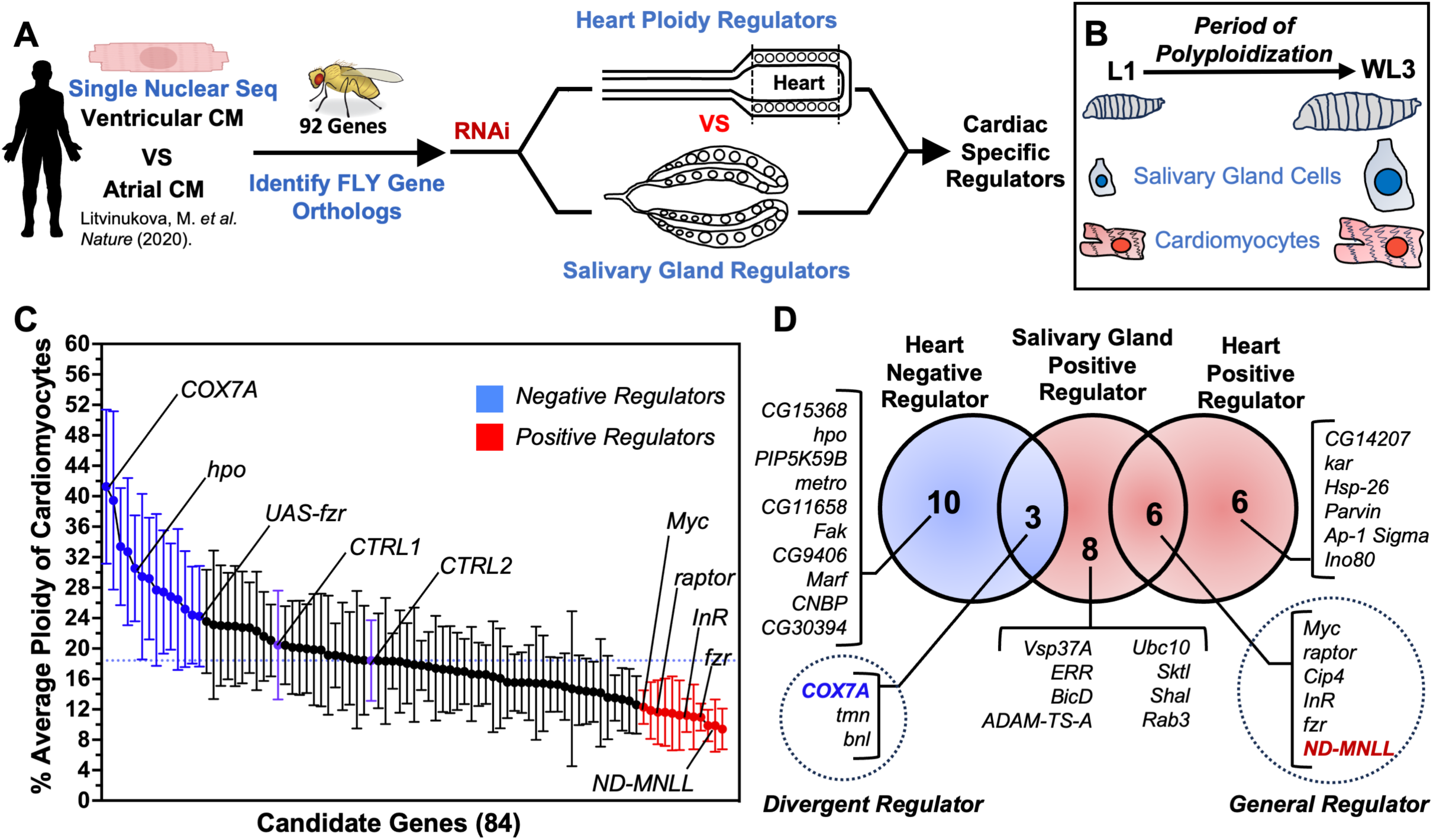
A reverse genetic screen of human ventricle enriched genes in *Drosophila* identifies tissue-specific ploidy regulation. **(A)** Schematic representation of the strategy used to identify cardiac-specific ploidy regulators. **(B)** Schematic of larval cardiomyocytes and salivary gland cells undergoing endoreplication during larval development to increase cell size. **(C)** Reverse genetic screen to identify ploidy regulators in *Drosophila* larval cardiomyocytes using orthologs of human cardiomyocyte expressed genes. CTRL1 and CTRL2 represent the two controls: *NP>V60000* and *NP>attP40*, respectively. Fzr Knockdown (*NP>fzr-RNAi)* and *Fzr* overexpression (*NP>UAS-fzr)* serve as positive controls for ploidy regulation. Mean±SD; For each genotype, 32 cardiomyocytes from two animals were analyzed to calculate the average ploidy (see Methods). **(D)** Venn diagram showing cardiac and salivary gland ploidy regulators. Red and blue color represents negative and positive ploidy regulators respectively. Dotted circles represent the Divergent ploidy regulators (3 genes: *COX7A*, tmn and *bnl*) and the General ploidy (6 genes: *Myc*, *Raptor*, *cip-4*, *InR*, *Fzr* and *ND-MNLL*).

Of the 92 genes tested (**TableS1: “Tab1-Ploidy Screen”**), 84 gene knockdowns produced viable WL3 (**FigS1A**), which we analyzed for cardiomyocyte ploidy changes in the heart chamber. We identified both positive (**Fig 1C, red**) and negative (**Fig 1C, blue**) regulators of cardiomyocyte ploidy. Since *fizzy-related (fzr) /Cdh1*, a well-established regulator and activator of the Anaphase-Promoting-Complex/Cyclosome (Cohen et al., 2018; Cohen et al., 2021; Schoenfelder et al., 2014; Sigrist and Lehner, 1997; Zielke et al., 2013), regulates larval cardiomyocyte polyploidization (Chakraborty et al., 2023), we used *NP>* to drive *UAS-fzr RNAi* and *UAS-fzr* overexpression as controls. Consistent with our previous finding, *NP>fzr RNAi* significantly reduces cardiomyocyte ploidy (Chakraborty et al., 2023), whereas *NP>*UAS-*fzr* increases cardiomyocyte ploidy (**Fig 1C**).

As our screen candidates were identified from human cardiomyocyte gene expression, we next assessed the specificity of each screen hit to cardiac ploidy. To do so, we took advantage of the fact that NP> simultaneously drives *UAS-RNAi* expression in another polyploid tissue, the salivary gland (**Fig 1A, 1B, S1B**). Our screen strain simultaneously expressed both *UAS-*RNAi constructs and a *UAS-mCherry-NLS* construct. *mCherry-NLS* expression is visible through the transparent larval cuticle, enabling us to quickly assay salivary gland size as a proxy of endoreplication (Zielke et al., 2011). Myc was included as a positive control, as it is a known endoreplication regulator that acts downstream of Fzr during salivary gland endoreplication (Qian et al., 2020). Indeed, *NP>UAS-myc RNAi* produces smaller glands (**FigS1F**). In total, we identified 17 positive regulators of salivary gland growth (**FigS1B-S)**. By comparing cardiomyocyte positive and negative ploidy regulators with salivary gland positive regulators (**Fig1D**), we identified three categories of genes: **1)** common regulators of polyploidization (core cell cycle regulators: *Myc*, *raptor*, *Cdc42-interacting protein 4* (*Cip4)*, Insulin-like receptor (*InR*), *fzr*, NADH dehydrogenase (ubiquinone) MNLL subunit (*ND-MNLL*); herein referred to as general ploidy regulators, **2)** genes that specifically regulate cardiomyocyte ploidy without affecting salivary gland ploidy, and vice versa; herein referred to as tissue-specific ploidy regulators, and **3)** genes with opposing effects on ploidy in the two organs, herein referred to as divergent ploidy regulators (Cytochrome c oxidase subunit 7A (*COX7A*), thinman (*tmn*), branchless (*bnl*)). This screen highlights new regulation of nuclear growth (ploidy) in the fly heart by genes that are enriched in the most highly polyploid human cardiomyocytes.

### COX7A negatively regulates cardiomyocyte polyploidization and heart organ growth

Notably, two genes that encode highly conserved orthologs of mitochondrial electron transport chain (ETC) components scored differently in our screen. *ND-MNLL*, a general ploidy regulator (**Fig 1D**, **Fig 2A, 2B**), encodes the MNLL subunit of NADH dehydrogenase (ubiquinone), a protein that is found by Cryo-EM in *Drosophila* mitochondrial respiratory chain complex I (Agip et al., 2023; Garcia et al., 2017). In contrast, *COX7A,* a divergent ploidy regulator (**Fig 1D**, **Fig 2A,2B, FigS2A,2B**), encodes *Cytochrome* c oxidase subunit 7A, a predicted subunit of mitochondrial respiratory chain complex IV. In the salivary gland, both *NP>ND-MNLL RNAi* and *NP>COX7A RNAi* decrease salivary gland ploidy (**Fig 2B, FigS2B**). *NP>ND-MNLL RNAi* knockdown hearts also exhibit decreased cardiomyocyte ploidy (**Fig 2A**), but *NP>COX7A RNAi* hearts instead show increased cardiomyocyte ploidy (**Fig 2A**). The distinct function of *COX7A* in ploidy regulation between these two tissues is mirrored in nuclear size-*NP>COX7A RNAi* cardiomyocytes (**Fig 2C vs. E**) have enlarged nuclei, while *NP>COX7A RNAi* salivary gland secretory cells have small nuclei (**Fig 2D vs. F**). A recent study of the zebrafish *COX7A* ortholog COX7A1 identified this protein as a key regulator of cardiac regeneration (García-Poyatos et al., 2024), but its role in ploidy, a common property of cardiomyocytes in many species (Hirose et al., 2019) has yet to be reported.

**Figure 2:**
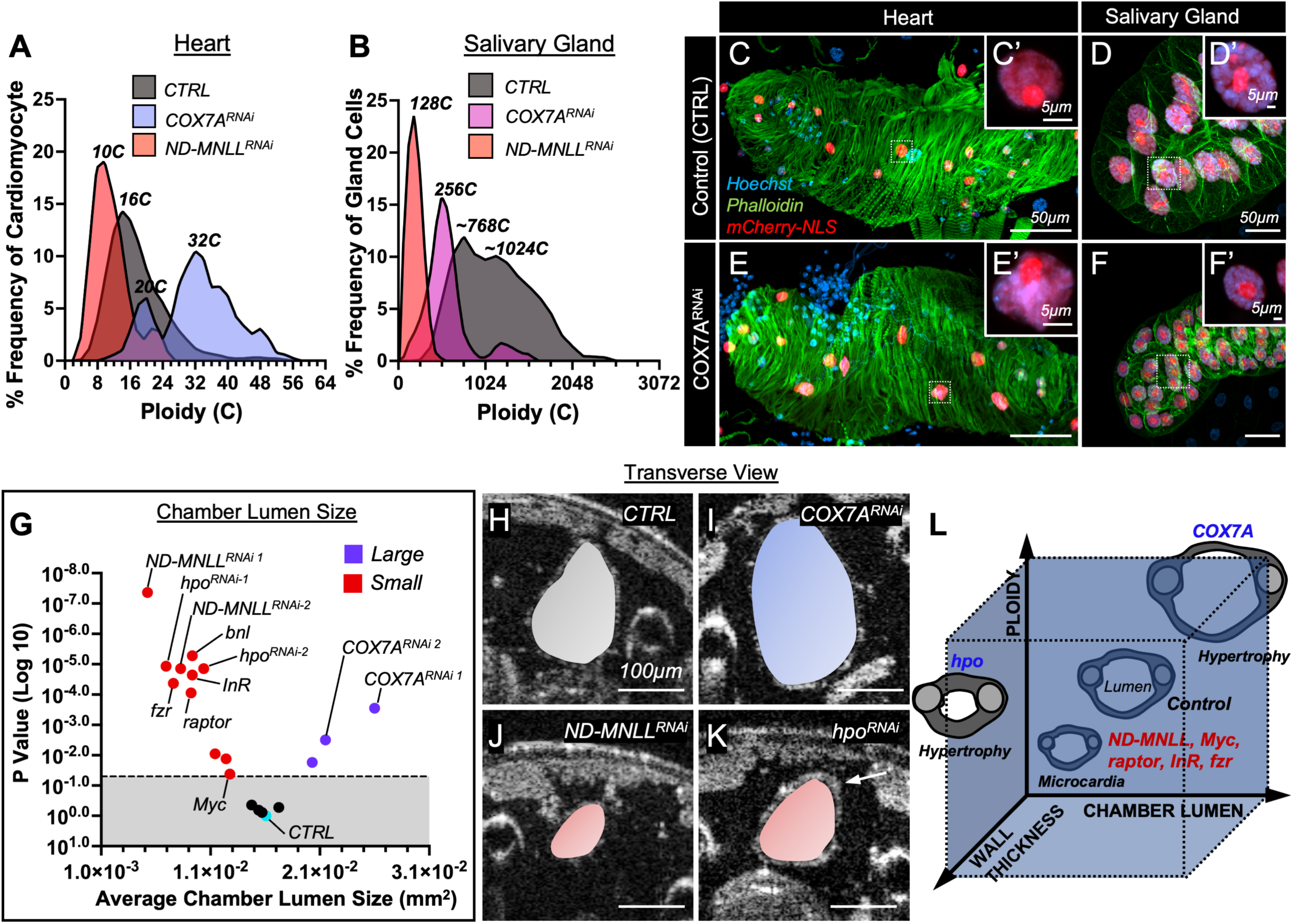
COX7A negatively regulates both tissue size and ploidy in the heart. **(A-B)** Ploidy distribution in Control (CTRL, *NP>V60000)*, *NP>COX7A RNAi* and *NP>MNLL RNAi* hearts (**A**) and Salivary glands (**B**). n>5 animals/group; For each animal, 16 cardiomyocytes in the heart chamber (A6–A5) and at least 20 posterior salivary gland cells were analyzed. e**(C-D’)** Representative images of Wandering Larvae 3 (WL3) heart (**C**) and salivary gland (**D**) from control (*NP>V60000*) animals. Insets show cardiomyocyte (**C’**) and gland cell (**D’**) nuclei. Chamber wall and gland membrane are labeled in green (Phalloidin staining), gland and cardiomyocyte nuclei in magenta (*NP>UAS-mCherry-NLS* and Hoechst), and all nuclei in blue (Hoechst). **(E-F’)** Representative images of WL3 heart (**E**) and salivary gland (**F**) from *NP>COX7A-RNAi* animals. Insets show cardiomyocyte (**E’**) and gland cell (**F’**) nuclei. Chamber wall and gland membrane are labeled in green (Phalloidin staining), gland and cardiomyocyte nuclei in magenta (*NP>UAS-mCherry-NLS* and Hoechst), and all nuclei in blue (Hoechst). **(G)** Volcano plot showing OCT measurements of End Diastolic Area (EDA, chamber lumen size) in WL3 heart chambers for heart ploidy regulators identified in the ploidy screen. Red and blue dots indicate gene knockdowns with significantly smaller or larger EDA, respectively (p < 0.05). Mean ± SD; unpaired two-tailed Student’s t-test. n ≥ 10 animals per group. **(H-K)** Representative transverse two-dimensional OCT images of the WL3 heart chamber showing EDA (pseudo-colored) for control (CTRL, *NP>V60000*) (B), *NP>COX7A-RNAi* (C), *NP>ND-MNLL RNAi* (D) and *NP>hpo RNAi* (E, white arrow indicates the chamber wall thickness). Each data set includes at least two biological repeats. **(L)** Schematic diagram illustrating the relationship between larval cardiomyocyte ploidy and chamber lumen size or wall thickness.

Ploidy correlates strongly with cell size (Morris et al., 2024). Previously, using Optical Coherent Tomography (OCT), we showed that cardiomyocyte cell size correlates with size of the cardiac chamber lumen area, and further that lumen area is significantly reduced in live WL3 *NP>fzr RNAi* or *NP>InR* RNAi animals, both of which have lower cardiomyocyte ploidy and microcardia (Chakraborty et al., 2023). Thus, we next examined the impact of COX7A and other selected ploidy screen hits on cardiac lumen size. To do so, we examined the transverse End Diastolic Area (EDA; chamber lumen size) of the heart chamber using OCT (see methods) (**Fig 2G; TableS1: “Tab2-PloidyScreenHits-OCT based”**). In multiple cases, both ploidy and lumen size increases (*RNAi* of *COX7A* and *CG15368*), resulting in cardiac hypertrophy (**Fig 1C**, **Fig 2G-I, L; TableS1: “Tab2-PloidyScreenHits-OCT based”**). Of note, our previous study found that constitutive activation of InR moderately increases heart ploidy (0.7-fold) but has no impact on chamber lumen size (Chakraborty et al., 2023). The magnitude of ploidy change for these new hypertrophic screen hits is higher than for activation of InR-approximately 2-fold (**Fig 1C**). This suggests that cardiac cell and organ size increases once ploidy crosses a certain threshold. We also found cases where both ploidy and chamber lumen size decrease (*RNAi* of *ND-MNLL*, *Myc*, *raptor*, *Hsp26* (Heat shock protein 26), *cip4*, *InR*, and *fzr*), resulting in microcardia (**Fig 1C**, **Fig 2J-L**). These results support the well-documented connection between ploidy (nuclear growth) and cell size (cytoplasmic growth) and highlight important regulators of optimal cell and organ size in the heart.

Not all tested genes impact ploidy and organ size in proportion. Specifically in *NP>hpo RNAi* or *NP>bnl RNAi* hearts, which impact Hippo and FGF signaling, respectively, cardiomyocyte ploidy increases while chamber lumen size decreases (**Fig 1C**, **Fig 2G,K**). In *NP>hpo RNAi* animals, this decrease in lumen size is accompanied by an increase in chamber wall thickness (**Fig 2K** (indicated by white arrow), **FigS2C**). Thus, *NP>hpo RNAi* animals exhibit a different class of hypertrophy whereby ploidy and cell thickness increase (**Fig 2L**) The ploidy and wall thickness *NP>hpo RNAi* phenotypes are consistent with previous findings on the role of Hippo signaling in the adult *Drosophila* heart (Yu et al., 2015). Our findings demonstrate a general correlation between ploidy (nuclear growth) and cardiac organ size (cytoplasmic growth), while suggesting that mechanical aspects of cardiac chamber shape (lumen size versus wall thickness) are coordinated by Hippo and FGF signaling (**Fig 2L**). Overall, our reverse genetic screen of human ventricle-enriched genes identified several regulators of nuclear and cytoplasmic growth and highlight COX7A as a negative regulator of ploidy and optimal organ size in the heart.

### Positive feedback between mitochondrial biogenesis and polyploidy during organ growth

COX7A family proteins are known to play a supportive function in mitochondrial Complex IV (García-Poyatos et al., 2024; Letts et al., 2016; Tsukihara et al., 1996; Wu et al., 2016), though the biology of COX7A proteins is complex, involving both tissue-and metabolism-dependent roles (Fernández-Vizarra et al., 2022; García-Poyatos et al., 2024; Sinkler et al., 2017). We therefore examined regulation of mitochondria by *Drosophila* COX7A, and its possible relation to ploidy and organ growth. We first examined mitochondrial size and density. To visualize mitochondria, we used the *UAS-mito-GFP* transgene (Pilling et al., 2006) driven by the *NP5169-Gal4* (*NP>*) driver. We compared mitochondrial size between control and *NP>COX7A RNAi* hearts in WL3 animals by measuring the area of individual mitochondria. While mitochondrial size remains unchanged in *NP>COX7A RNAi* hearts (Fig 3A**-A’,C-C’, FigS3A**), it decreases in *NP>COX7A RNAi* salivary glands (**Fig 3B-B’,D-D’, FigS3B**). We next measured cellular mitochondrial density (see methods). *NP>COX7A RNAi* hearts, but not salivary glands, display a marked increase in mitochondrial density (**Fig 3E vs FigS3C**). This mitochondrial phenotype is consistent with previous studies of COX7A orthologs in mice and zebrafish (García-Poyatos et al., 2024; Huttemann et al., 2012). Overall, our findings suggest that COX7A represses mitochondrial biogenesis specifically in the heart, and that this increased mitochondrial density correlates with increased heart organ size (cytoplasmic growth) and cardiomyocyte ploidy (nuclear growth).

**Figure 3:**
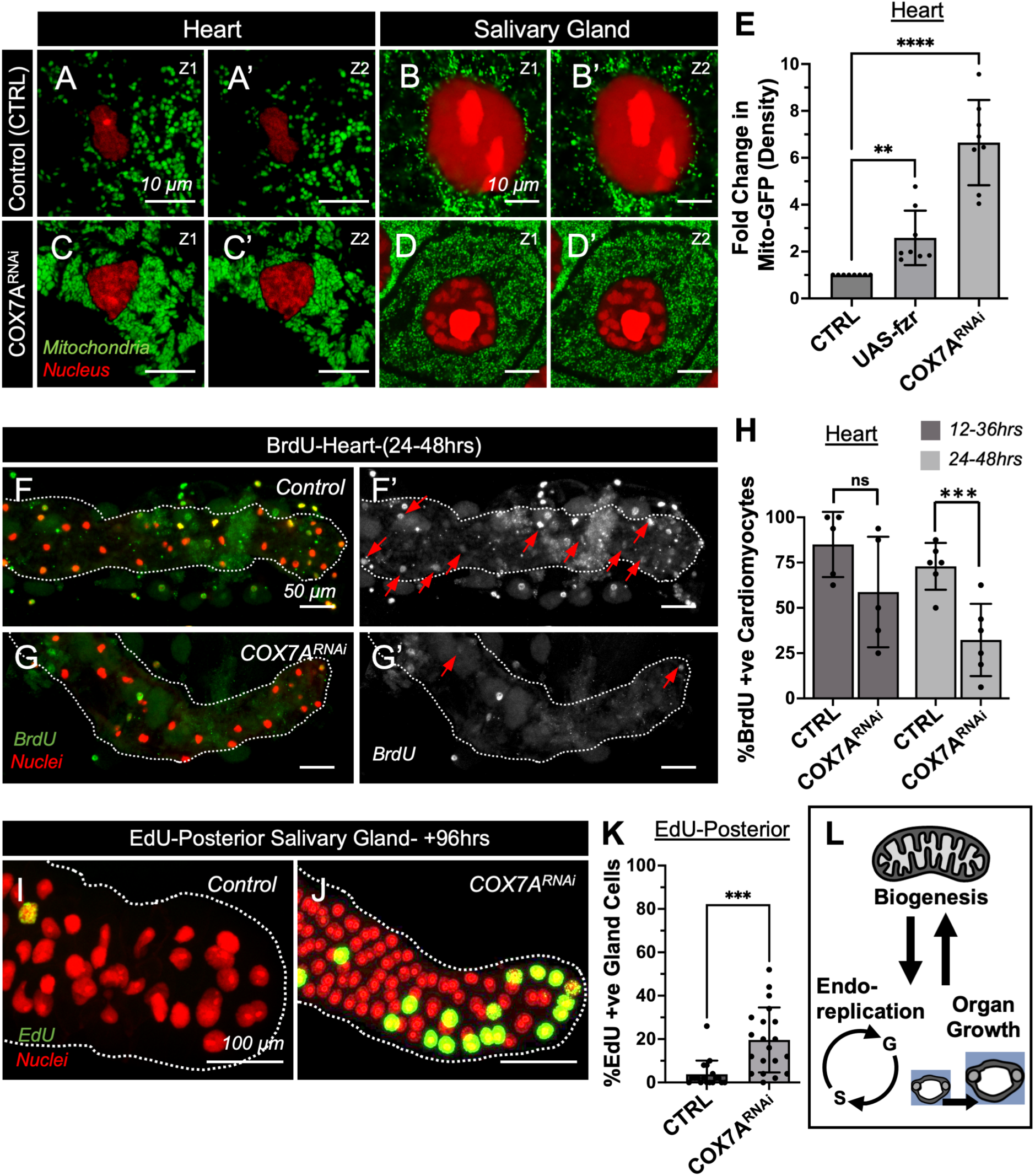
C**O**X7A **coordinates tissue-specific mitochondrial biogenesis, endoreplication and organ growth. (A–B**′) Single Z-section images (Z1 and Z2) of representative Wandering Third Instar Larvae (WL3) heart (A, A′) and salivary gland (B, B′) from control animals (*NP>V60000*). Mitochondria are labeled in green using *NP>mito-GFP*. Cardiomyocyte nuclei (**A, A**′) and gland cell nuclei (**B, B**′) are labeled in red using *NP>UAS-mCherry-NLS*. **(C–D**′) Single Z-section images (Z1 and Z2) of representative WL3 heart (**C, C**′) and salivary gland (**D, D**′) from *NP>COX7A-RNAi* animals. Mitochondria are labeled in green using *NP>mito-GFP*. Cardiomyocyte nuclei (**C, C**′) and gland cell nuclei (**D, D**′) are labeled in red using *NP>UAS-mCherry-NLS*. **(E)** Fold change in fluorescence intensity of *NP>UAS-mito-GFP* labeled mitochondria in the heart chamber for the indicated genotypes. Mean±SD; **P < 0.001, ****P <0.0001; Unpaired two-tailed Student’s t-test. n ≥ 5 animals per group. Larval heart segment A7-A5 were imaged. **(F–G**′) Representative images of BrdU-positive cardiomyocytes (**F, G**: green; **F**′**, G**′: white) in the WL3 heart chamber of control (**F, F**′; *NP>V60000*) and *NP>COX7A RNAi* (**G, G**′) animals. Cardiomyocyte nuclei are labeled in red (*NP>UAS-mCherry-NLS*). Animals were pulse-fed BrdU for 24 hours between 24–48 hours post-hatching (*see Methods*). Red arrows (**F**′**, G**′) indicate BrdU-positive cardiomyocytes. **(H)** Percentage of BrdU-positive cardiomyocytes in the heart chamber at the indicated time points after hatching. Mean ± SD; ^ns^P>0.05, **P < 0.001; Unpaired two-tailed Student’s t-test. n ≥ 5 animals per group. Experiments were repeated at least three times. **(I,J** Representative images of EdU-positive posterior gland cells (**I,J**: green) in the WL3 salivary glands of control (I; *NP>V60000*) and *NP>COX7A RNAi* (**J**) animals. Gland cell nuclei are labeled in red (*NP>UAS-mCherry-NLS*). WL3 salivary glands were soaked in EdU for 2 hours prior to fixation (see Methods). **(K)** Percentage of EdU-positive posterior gland cells. Mean ± SD; **P < 0.001; Unpaired two-tailed Student’s t-test. n ≥ 5 animals per group. Each data set includes at least two biological repeats. **(L)** Schematic diagram illustrating the reciprocal relationship between mitochondrial biogenesis, endoreplication, and organ growth.

Given the substantial impact on ploidy by *COX7A RNAi* (**Fig 1C, D, Fig 2A-F**), we next examined how COX7A affects DNA replication dynamics during endoreplication in the heart and salivary gland. We used pulses of thymidine analogs (BrdU or EdU) to assay replication (see methods). In our previous study, we showed that cardiomyocytes in control hearts undergo endoreplication between 0-48 hours post-hatching (Chakraborty et al., 2023). Consistent with those findings, approximately 85% and 72% of cardiomyocytes in control hearts are BrdU-positive following a 24-hour BrdU pulse during either the 0-24 or 24-48 hour post-hatching windows, respectively (**Fig 3F,F**′**,H**). In contrast, *NP>COX7A RNAi* hearts show a significant reduction in BrdU-positive cardiomyocytes, with only 58.7% and ∼32% labeled during the same respective time windows (**Fig 3G-H**). These results, together with the increased cardiomyocyte ploidy observed in *NP>COX7A RNAi* hearts, suggest that COX7A knockdown shortens the time spent in each endoreplication S-phase, enabling cardiomyocytes to reach higher ploidy levels more rapidly.

Next, we examined endoreplication in salivary glands. Salivary gland endoreplication spans approximately 7-96 hours after egg deposition (Zielke et al., 2011). We examined DNA replication near the end of this window, by performing a 2-hour EdU soak assay in dissected WL3 salivary glands (see Methods) at 96 hours post-hatching. Since posterior gland secretory cells are larger and more polyploid than anterior cells (**FigS1B**, anterior to the left), we quantified EdU-positive nuclei only in the posterior region. Consistently, compared to controls, *NP>COX7A RNAi* salivary glands show an increased number of EdU-positive nuclei specifically in the posterior region (**Fig 3I-K**). Taken together with the decreased ploidy (**Fig 2B**), these results suggest that *NP>COX7A RNAi* salivary glands exhibit prolonged time in each endoreplication S-phase. Together, these results suggest that the differential regulation of mitochondrial biogenesis by COX7A in the heart and salivary gland reflects its tissue-specific roles in controlling DNA replication (ploidy), and organ size (**Fig 3L)**.

Our results with COX7A suggest that increased mitochondrial biogenesis in the cytoplasm drives increased nuclear ploidy in the heart. We next tested whether the converse is true-that increased nuclear ploidy may drive increased mitochondrial biogenesis in the cytoplasm. Since overexpression of the endoreplication regulator *fzr* increases ploidy (**Fig 1C**), we examined mitochondrial density in the heart and salivary gland of these animals. Indeed, *NP>UAS-fzr* animals exhibit increased mitochondrial density in both hearts and salivary glands (**Fig 3E, S3C**). Overall, our findings highlight positive feedback between mitochondrial biogenesis and the rate of endoreplication and suggest that COX7A is critical for achieving optimal organ growth.

### COX7A’s cardiac growth repressor function is distinct among ETC genes

Our findings with COX7A suggests that the same gene positively promotes mitochondrial biogenesis and ploidy in one tissue (salivary gland) represses these processes in another tissue (heart). Previous work in mouse decidual cells suggests that electron transport chain (ETC) components positively regulate ploidy through mitochondrial biogenesis (Ma et al., 2011), similar to our findings here with *ND-MNLL* in the salivary gland and heart and with *COX7A* in the salivary gland (but not heart) (**Fig 2A,B**). Thus, our results highlight potential differences in annotated ETC gene function between tissues with regards to mitochondrial and ploidy regulation.

To further investigate whether other electron transport genes might function similarly to *COX7A* in the heart, we conducted an OCT-based screen targeting 55 ETC genes, including 7 mitochondrially encoded and 48 nuclear encoded ETC genes (**Fig 4A-A’; TableS1: “Tab3-ETC Screen-OCT based”**). We used lumen size (transverse End Diastolic Area (EDA)) as a proxy for ploidy, given the correlation between these two measurements seen in our other ETC knockdowns, namely *NP>COX7A RNAi* and *NP>MNLL* (Fig 2G). Among 26 gene knockdowns that show significant lumen size changes (**Fig 4B-F, FigS4A-F)**, only *NP>COX7A RNAi* increases cardiac lumen size, whereas the other 25 reduce it (**Fig 4A’**). In contrast, all 26 gene knockdowns (*NP>COX7A RNAi* included) decrease salivary gland size (**Fig 4F**). These findings indicate that, among 55 ETC genes tested, only *COX7A* has a unique role as a repressor of nuclear (ploidy) and cytoplasmic (lumen size) growth in the heart.

**Figure 4:**
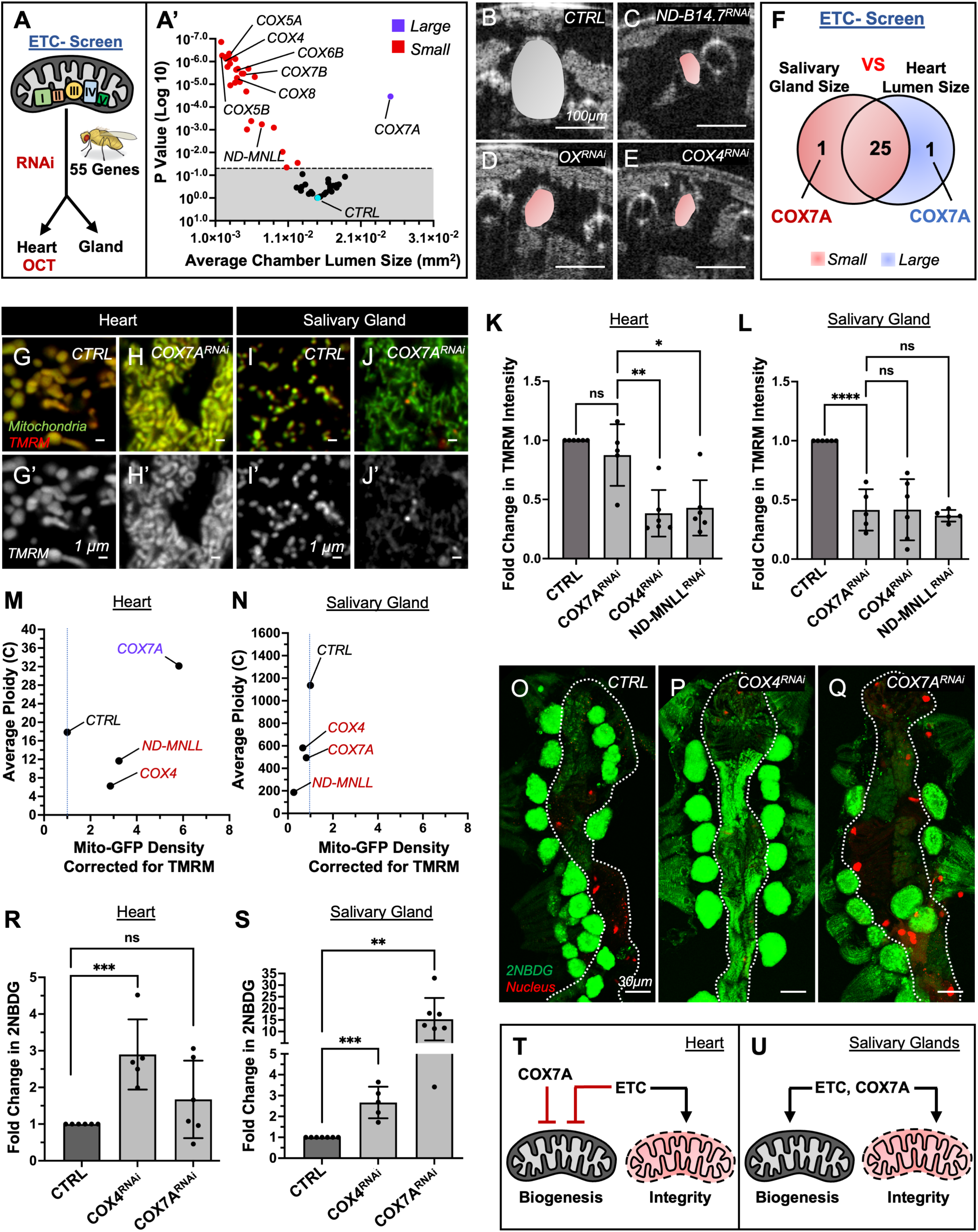
COX7A is required for mitochondrial integrity in salivary glands but not in hearts. **(A)** Schematic illustrating the experimental approach used to identify ETC genes that regulate heart chamber lumen size and salivary gland size. **(A**′) Volcano plot showing OCT measurements of end-diastolic area (EDA, Chamber lumen size) in WL3 heart chambers for 55 electron transport chain (ETC) gene knockdowns. Red and blue dots indicate gene knockdowns that significantly decrease or increase EDA, respectively (p < 0.05). Mean ± SD; Unpaired two-tailed Student’s t-test. n ≥ 5 animals per group. Mean ± SD; Unpaired two-tailed Student’s t-test. **(B–E)** Representative transverse two-dimensional OCT images of WL3 heart chambers showing EDA (pseudo-colored) for control (**B**; Control (CTRL), *NP>V60000*), *NP>ND-B14.7 RNAi* (**C**; Complex I), *NP>ox RNAi* (**D**; Complex III), and *NP>COX4 RNAi* (**E**; Complex IV). **(F)** Venn diagram showing the overlap between heart chamber lumen size and salivary gland size phenotypes for ETC gene knockdowns. **(G–J**′) Representative images of TMRM-stained mitochondria in control (**G, G**′**, I, I**′) and *NP>COX7A RNAi* (**H, H**′**, J, J**′) hearts (**G–H**′) and salivary glands (**I–J**′). Mitochondria are co-labeled with *NP>UAS-mito-GFP* (green, **G–J**) and TMRM (red, **G–J**; white, **G**′**–J**′). **(K–L)** Fold change in TMRM intensity for the indicated genotypes in the heart chamber (**K**) and salivary gland (**L**). Mean ± SD; ^ns^P>0.05, ** P < 0.01, **** P < 0.0001; Unpaired two-tailed Student’s t-test. n ≥ 5 animals per group. Each data set includes at least two biological repeats. **(M–N)** Scatter plots showing total mitochondrial GFP fluorescence density (intensity/area) for functional mitochondria versus average ploidy for the indicated genotypes in the heart chamber (**M**) and salivary glands (**N**). n ≥ 6 animals per group. Each data set includes at least two biological repeats. **(O–Q)** Representative images of 2-NBDG-labeled (green) heart chambers for the indicated genotypes. **(R–S)** Fold change in 2-NBDG intensity for the indicated genotypes in the heart chamber (**R**) and salivary gland (**S**). Mean ± SD; ^ns^P>0.05, ** P < 0.01, *** P < 0.001; Unpaired two-tailed Student’s t-test. n ≥ 5 animals per group. Each data set includes at least two biological repeats. **(T–U)** Schematic diagrams showing the regulation of ETC genes and COX7A in mitochondrial biogenesis and integrity in the heart (**T**) and salivary glands (**U**).

We next examined mitochondria in selected ETC gene RNAi lines. Similar to *COX7A RNAi*, knockdown of other ETC genes including *ND-MNLL* and the annotated Complex IV subunit gene *COX4* drastically increase mitochondrial density in the heart (**FigS4G**), but not in salivary glands (**FigS4H**). These findings suggest that compromising function of annotated ETC genes triggers mitochondrial biogenesis specifically in the heart. However, this finding does not match our observed positive feedback between mitochondrial biogenesis and endoreplication (**Fig 3L**). To uncover clues as to the disconnect between increased mitochondria but decreased ploidy in all of our ETC gene knockdowns except COX7A, we examined heart function. Many ETC gene knockdowns, including ploidy-reduced *NP>ND-MNLL RNAi* hearts, but not *NP>COX7A RNAi* hearts, exhibit reduced heartbeat. (**FigS4I; TableS1: “Tab4-Heartbeat-OCT based”**). Mitochondrial dysfunction is known to impair ATP production and cause contractile failure (Zhou and Tian, 2018). We thus hypothesized that preserved mitochondrial integrity may underlie the uniquely increased ploidy in *NP>COX7A RNAi* hearts. Mitochondrial membrane integrity is a readout of mitochondrial function (Schenkel and Bakovic, 2014). Therefore, to test mitochondrial integrity, we assessed membrane potential using TMRM (Tetramethylrhodamine, methyl ester) staining (**FigS4J**). TMRM is a membrane-permeable dye that accumulates in the mitochondrial matrix of active mitochondria in living cells, whereas dysfunctional mitochondria fail to sequester the dye (Perry et al., 2011; Scaduto and Grotyohann, 1999). We compared mitochondrial membrane potential in *NP>COX7A RNAi* to other ETC gene knockdowns (*ND-MNLL*-complex I and *COX4*-annotated complex IV) in hearts and salivary glands. Indeed, in *NP>COX7A RNAi* hearts, mitochondrial membrane potential is unaffected (**Fig 4G-H’, K**), whereas it is significantly reduced in *NP>COX7A RNAi* salivary glands (**Fig 4I-J’,L**). By contrast, *NP>MNLL RNAi* and *NP>COX4 RNAi* causes a marked reduction in mitochondrial membrane potential in both hearts (**Fig 4K, FigS4K,M,N**) and salivary glands (**Fig 4L, FigS4K’,M’,N’**). These results reveal that COX7A functions similarly to other ETC components to regulate mitochondrial integrity in the salivary gland. However, in the heart, COX7A lacks a role in mitochondrial integrity.

Previous studies have shown that blockage of the electron transport chain (ETC) forces cells to rely on anaerobic respiration by upregulating glucose uptake (Wu et al., 2024). This phenomenon can be measured in living cells by quantifying the uptake of fluorescently labeled glucose, 2NBDG (2-(N-(7-Nitrobenz-2-oxa-1,3-diazol-4-yl)Amino)-2-Deoxyglucose), which becomes trapped inside the cell after uptake because it cannot be further metabolized through glycolysis (**FigS4Q**) (O’Neil et al., 2005; Wong et al., 2019). Consistent with our TMRM data, 2NBDG uptake is significantly increased in *NP>COX4 RNAi* hearts (**Fig 4O,P,R**) and salivary glands (**Fig4S, FigS4R,T**). Similarly, 2NBDG uptake is significantly upregulated in *NP>COX7A RNAi* salivary glands (**Fig4S, FigS4U**). However, 2NBDG uptake is not significantly upregulated in *COX7A* knockdown hearts (**Fig 4Q,R**). Taken together, we observe that when mitochondrial integrity is compromised as a result of ETC component knockdown (except COX7A), cells of the heart and salivary gland switch to glycolysis, and ploidy is reduced (**Fig 4M,N,T**). The most likely explanation of the ploidy reduction is that it is a secondary consequence of compromised mitochondria. To test if ploidy reduction itself is sufficient to switch cells to glycolysis, we examined *NP>fzr RNAi* hearts and salivary glands. Neither 2NBDG uptake (**FigS4S,V,W**) nor TMRM fluorescence (**FigS4L,L’,O,P)** is impacted by *NP>fzr RNAi* in these tissues. Thus, reducing ploidy is not sufficient to switch to glycolysis.

Overall, we find that, among all annotated ETC genes, only COX7A shows distinct functions in the heart versus the salivary gland, with its cardiac role independent of mitochondrial integrity (**Fig 4T,U**). This lack of regulation of mitochondrial integrity, coupled with a role in repressing mitochondrial biogenesis, provides an explanation for the unique role of cardiac COX7A in repression of cytoplasmic growth through mitochondrial biogenesis and nuclear growth (ploidy).

### Cardiac ploidy repression by COX7A is associated with its mitochondrial isoform

Our results suggest a non-ETC function of COX7A in the heart to control mitochondrial biogenesis and cardiac polyploidy. We next explored the underlying molecular mechanism, particularly whether COX7A primarily functions in the mitochondria. The *Drosophila COX7A* gene has two isoforms, A and B, which differ in their 5’ UTR and transcription start sites but share a similar 3’ UTR and common cytochrome c oxidase (COX) domain (**Fig 5A,B**). Interestingly, analysis with TargetP-2.0 (Almagro Armenteros et al., 2019), reveals that only isoform A contains a mitochondrial targeting signal (**Fig 5A-B**). Using isoform-specific primers (methods), we find that both COX7A isoforms are significantly reduced by *NP>COX7A RNAi* in the salivary gland (**FigS5A,B**) and the heart (**FigS5C,D**). To examine the role of each *COX7A* isoform separately, we generated *UAS-GFP*-tagged, RNAi-resistant transgenes for each (methods). When the transgenes are expressed in S2 cells, the MTS-containing isoform A colocalizes with mitochondria (**FigS5E,E’**), whereas isoform B primarily localizes to the nucleus (**FigS5F,F’**). We then generated transgenic flies expressing each *COX7A* isoform transgene. As in S2 cells, isoform A localizes to mitochondria in both heart and salivary glands (**Fig 5C,D**), while isoform B localizes to the nucleus in both organs (**Fig 5E,F**).

**Figure 5:**
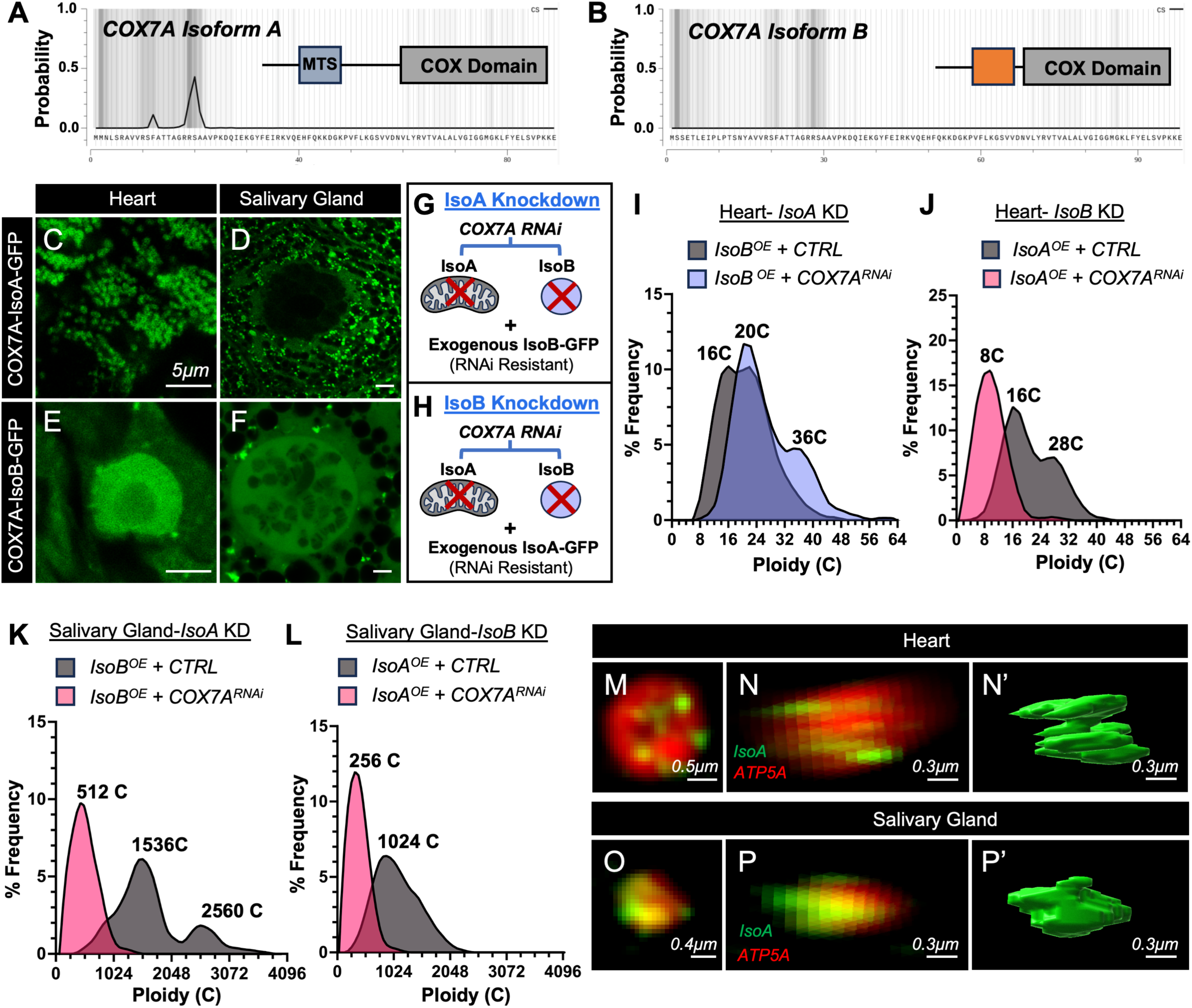
The mitochondrial COX7A Isoform negatively regulates cardiac ploidy, unlike in salivary glands. **(A-B)** TargetP-2.0 analysis of COX7A Isoform A (**A**) and Isoform B (**B**), as described in the Methods section. **(C-D)** Representative images of the heart chamber (**C**) and salivary gland (**D**) showing mitochondrial localization of overexpressed GFP-tagged COX7A Isoform A. **(E-F)** Representative images of the heart chamber (**E**) and salivary gland (**F**) showing nuclear localization of overexpressed GFP-tagged COX7A Isoform B. **(G-H)** Schematics illustrating the knockdown strategy for COX7A Isoform A (**G**) and Isoform B (**H**). **(I-J)** Ploidy distribution in Control and *NP>COX7A-RNAi* hearts with overexpression of RNAi resistant Isoform B (**I**) and Isoform A (**J**). n>5 animals/group. For each animal, the ploidy of 16 cardiomyocytes in the heart chamber (A6–A5) was analyzed. Each data set includes at least two biological repeats. **(K-L)** Ploidy distribution in Control and *NP>COX7A-RNAi* salivary glands with overexpression of RNAi resistant Isoform B (**K**) and Isoform A (**L**). n>5 animals/group. For each animal, at least 20 posterior salivary gland cells were analyzed. Each data set includes at least two biological repeats. **(M-P’)** Super-resolution imaging of mitochondria in heart chamber (**M-N’**) and salivary gland (**O-P’**) expressing GFP tagged COX7A-Isoform A (Green). Mitochondria are stained with ATP5A antibody (red). Top View (**M, O**) and lateral view (**N, P**). 3D-rendered images of GFP tagged COX7A-Isoform A (Green) in heart (**N’**) and salivary gland (**P’**).

To next determine the impact of each COX7A isoform on ploidy, we performed rescue experiments by individually overexpressing *UAS-GFP-COX7A-*isoform A and *UAS-GFP-COX7A-*isoform B in a *NP>COX7A RNAi* background (**Fig 5G,H**). The RNAi-resistant nature of the constructs enabled us to therefore study Isoform A knockdown (*NP>UAS-GFP-COX7A-*isoform B + *NP>COX7A RNAi*) and Isoform B knockdown (*NP>UAS-GFP-COX7A-*isoform A + *NP>COX7A RNAi*) (**Fig 5 G,H**). Mitochondrial *IsoA* knockdown increases larval heart ploidy (**Fig 5I**) but decreases salivary gland ploidy (**Fig 5K**), consistent with our *NP>COX7A RNAi* results (**Fig 2A-F**). In contrast, nuclear *IsoB* knockdown decreases ploidy in both larval hearts (**Fig 5J**) and salivary glands (**Fig 5L**). Taken together with our COX7A whole gene knockdown results, these isoform-specific results highlight distinct roles of each COX7A isoform in cardiomyocyte ploidy regulation, while suggesting that the cardiac ploidy repression phenotype seen in *NP>COX7A RNAi* is primarily due to isoform A function. Consistent with our data that COX7A functions distinctly in the heart and salivary gland to regulate mitochondrial integrity, super-resolution microscopy and 3D rendering (see methods) shows that UAS-GFP-tagged isoform A structures in heart mitochondria (**Fig 5M-N’**) appear different from those in salivary glands (**Fig 5O-P’**). Together, these results suggest that the unique role of COX7A in repressing cardiac polyploidy is through mitochondrial isoform A.

### Human COX7A1 represses cardiomyocyte ploidy in iPSC-CMs

COX7A family members are linked to heart organ size in both zebrafish and mice (García-Poyatos et al., 2024; Huttemann et al., 2012). Our findings from cladogram analysis of annotated mitochondrial respiratory chain complex IV components (methods) suggest that COX7A family members are only present in organisms with hearts (**Fig 6A**), perhaps consistent with the emergence of COX7A as an important regulator of ploidy-based organ growth that plays a distinct role in the heart. Given our finding of distinct functions of mitochondrial and nuclear COX7A gene products in the *Drosophila* heart, we next examined evolutionary divergence of the COX7A gene family. To do so, we performed phylogenetic analysis of the 5’ UTRs of COX7A isoforms in *Drosophila* and several model vertebrates (humans, rat, mouse, and zebrafish), based on TargetP-2.0 predictions. Isoforms with higher or lower probability of containing a mitochondrial targeting sequence were analyzed together (See Methods, **TableS2**). Interestingly, Simple Phylogeny (Madeira et al., 2024) analysis revealed that the mitochondrial *Drosophila* isoform A and the nuclear isoform B are closely related to human *COX7A1* (**Fig 6B**) and human *COX7A2L* isoform B (**Fig 6C**), respectively. Therefore, fly COX7A isoform A resembles human COX7A1 in terms of containing a mitochondrial targeting sequence.

**Figure 6:**
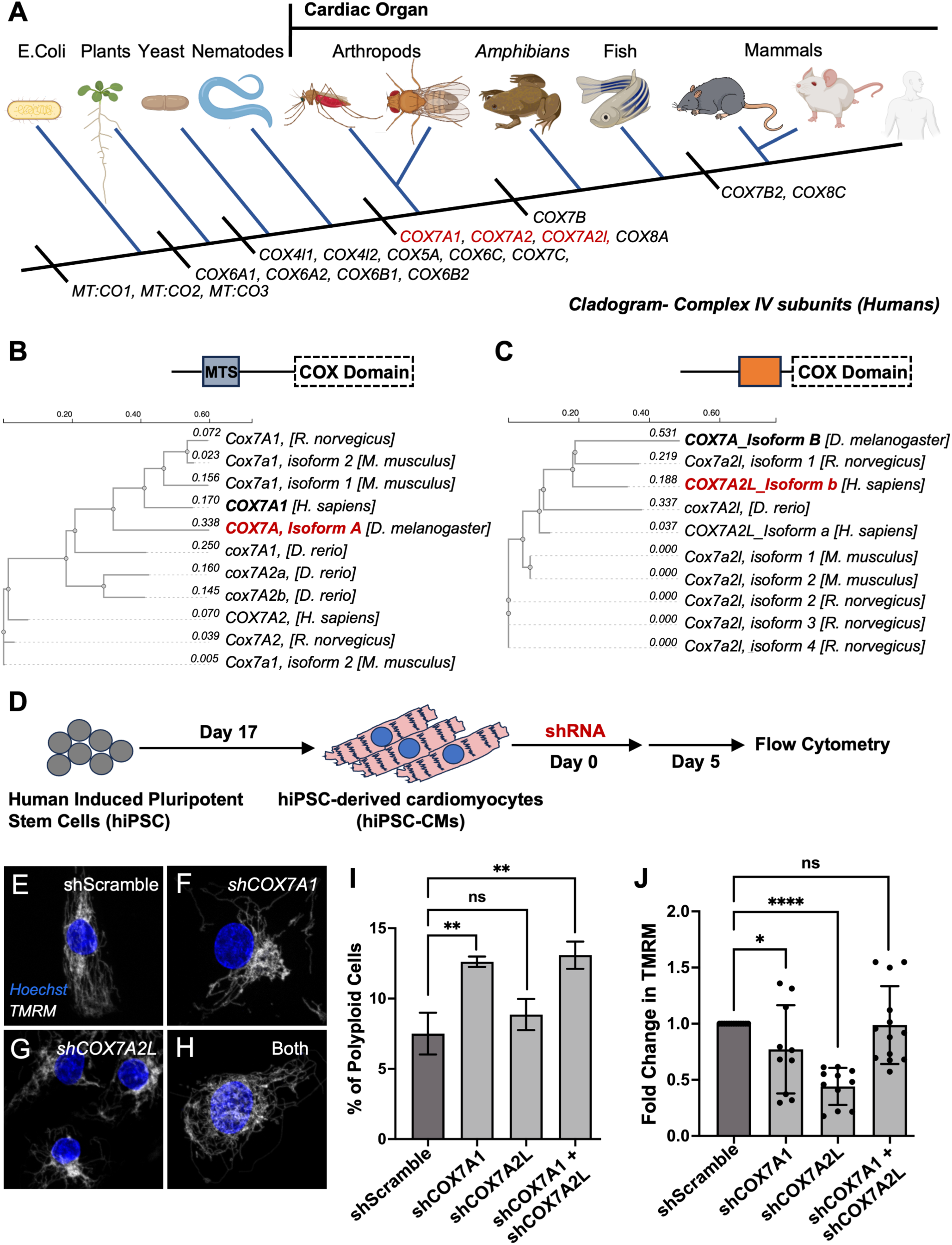
C**O**X7A1 **evolved in heart-bearing species and suppress cardiomyocyte ploidy in hiPSC-CMs (A)** Evolutionary relationship of mammalian Complex IV subunits (Cladogram analysis; see Methods), highlighting the emergence of COX7A genes in organisms with a developed heart. **(B-C)** Phylogenetic analysis of TargetP-2.0 analyzed COX7A orthologs with high-probability mitochondrial targeting signals (**B**) and without such signals (**C**). The COX domain was excluded from the analysis (see Methods). **(D)** Schematic representation of the strategy used to knock down orthologous COX7A isoforms in human iPSC-derived cardiomyocytes. **(E-H)** Representative images of control (E, shScramble), shCOX7A1 (**F**), sh-COX7A2L (**G**), and combined shCOX7A1 & shCOX7A2L (**H**) in human iPSC-derived cardiomyocytes. Nuclei are stained with Hoechst, and mitochondria with TMRM. **(I)** Percentage of polyploid cells in hiPSC-CMs following shRNA-mediated knockdown of the indicated genes, as measured by flow cytometry. Mean±SD; ****P <0.0001, ***P<0.001, ^ns^P>0.05; Unpaired two-tailed Student’s t-test. Each data set includes at least two biological repeats. **(J)** Fold change in TMRM intensity in hiPSC-CMs following shRNA-mediated knockdown of the indicated genes. Mean ± SD; ^ns^P>0.05, ** P < 0.01, **** P < 0.0001; Unpaired two-tailed Student’s t-test. 10 cells were analyzed from each group.

Given our findings of a role for fly *COX7A* isoform A in cardiac ploidy repression, we next tested whether *COX7A1* similarly regulates cardiac ploidy in humans. We thus knocked down *COX7A1* in human iPSC-derived cardiomyocytes (hiPSC-CMs) using shRNA (**Fig 6D, FigS6A,A’**). Knockdown efficiency of *COX7A1* and *COX7A2L* was confirmed by RT-qPCR (**FigS6B,C**). *COX7A1* knockdown consistently increases polyploidy in cardiomyocytes, as measured by flow cytometry (**Fig 6E,F,I**). By contrast, shRNA-mediated knockdown of *COX7A2L* in hiPSC-CMs does not increase cardiomyocyte polyploidization (Fig 6G,I). Next, we examined mitochondrial integrity. Double knockdown of human *COX7A1* and *COX7A2L* (analogous to fly *COX7A RNAi*) preserves mitochondrial integrity (Fig 6J), as we observe in flies. Further, individual depletion of each gene alone shows that COX7A2L is mainly required for mitochondrial integrity compared to COX7A1 (Fig 6J). Overall, our results indicate that in both flies and human cells, COX7A gene products containing a mitochondrial targeting sequence function to repress cardiac polyploidy.

### COX7A is required for normal heart function

Our findings reveal a differential mitochondrial regulatory network between salivary glands and hearts. Specifically, we find that, among all annotated ETC genes, only cardiac COX7A lacks canonical ETC control of mitochondrial integrity yet retains a role in mitochondrial biogenesis. We next tested the role of mitochondrial integrity in COX7A cardiac ploidy repression. To do so, perturbed mitochondrial integrity independently of core ETC regulation. We thus examined the Prohibitin 1 (*Phb1*) gene. *Phb1* and the related *Phb2* are known regulators of mitochondrial membrane integrity that function outside of the ETC (Oyang et al., 2022; Signorile et al., 2019; Wei et al., 2017). *NP>Phb1 RNAi* compromises mitochondrial integrity (**Fig 7C, FigS7C**) and density of functional mitochondria in both the heart and salivary gland (**FigS7D, FigS7D**). Consistent with our finding of a requirement for mitochondrial integrity in achieving optimal ploidy and organ size, both *NP>Phb1 RNAi* and *NP>Phb2 RNAi* hearts show significantly reduced ploidy (**Fig 7E, FigS7E**) and heart chamber lumen size (**Fig 7F-H,J**). These findings enabled us to use *Phb1* depletion in a *COX7A* knockdown background to test if mitochondrial integrity is required for COX7A ploidy repression in heart. We thus performed double knockdowns of *Phb1* and *COX7A*. Indeed, reduction of *Phb1* in *NP>COX7A RNAi* hearts decreases mitochondrial integrity (**Fig 7A-C**), density (**Fig 7D**) and cardiomyocyte ploidy (**Fig 7D,E**).

**Figure 7:**
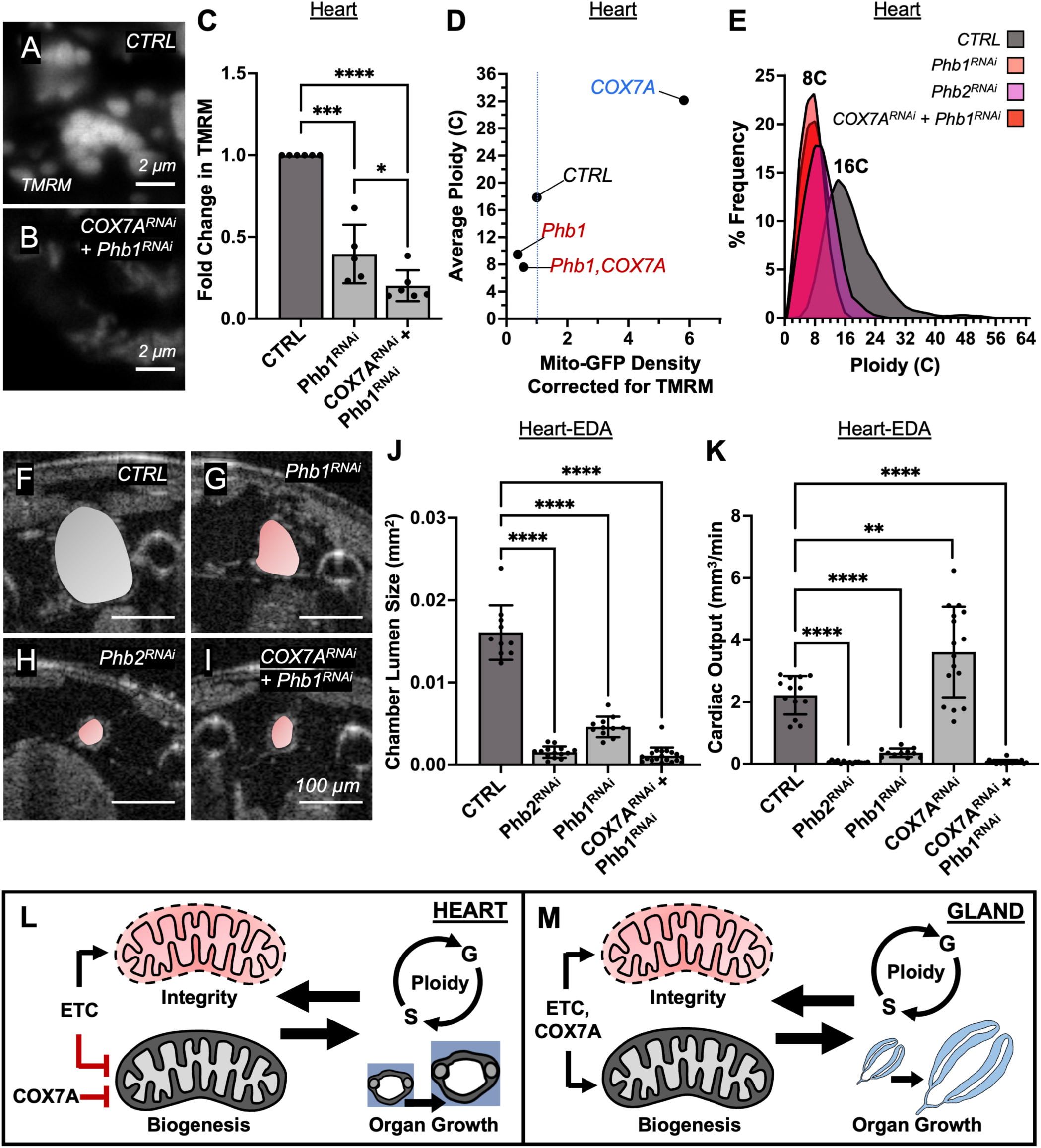
COX7A-mediated repression of cardiac ploidy requires mitochondrial function. **(A–B)** Representative images of TMRM-stained mitochondria in the heart chamber of control (A; *NP>V60000*) and *NP>Phb1-RNAi;COX7A-RNAi* (**B**) animals. **(C)** Fold change in TMRM intensity in the heart chamber for the indicated genotypes. Mean±SD; ^ns^P>0.05, *P < 0.05, ***P < 0.01, ****P < 0.0001; Unpaired two-tailed Student’s t-test. n ≥ 5 animals per group. Each data set includes at least two biological repeats. **(D)** Scatter plots showing total mitochondrial GFP fluorescence density (intensity/area) for functional mitochondria versus average ploidy for the indicated genotypes in the heart chamber. n ≥ 5 animals per group. **(E)** Ploidy distribution in the heart chamber of indicated genotypes. n>5 animals/group. For each animal, the ploidy of 16 cardiomyocytes in the heart chamber (A6–A5) were analyzed. Each data set includes at least two biological repeats. **(F-I)** Representative transverse two-dimensional OCT images of WL3 heart chambers showing EDA (pseudo-colored) for control (**F**; CTRL, *NP>V60000*), *NP>Phb1-RNAi* (**G**), *NP>Phb2-RNAi* (**H**), and *NP>Phb1-RNAi;COX7A-RNAi* (**I**). **(J)** Quantitative analysis of heart chamber lumen size (EDA) in WL3 heart chambers for the indicated genotypes. Mean±SD; ***P < 0.0001; unpaired two-tailed Student’s t-test. n ≥ 10 animals per group. Each data set includes at least two biological repeats. **(L-M)** Schematic diagrams of the proposed model for COX7A isoform A-mediated regulation of tissue-specific endoreplication.

Finally, we examined the role of COX7A in heart lumen size and function, and the dependency of these important cardiac metrics on mitochondrial integrity. We first examined chamber lumen size. *NP>COX7A RNAi* hearts exhibit increased chamber lumen size, indicative of cardiac hypertrophy (**Fig 2G-I,L, Video S1**). This hypertrophy depends on mitochondrial integrity, as double *NP>Phb1 RNAi;COX7A RNAi* drastically reduces chamber lumen size, indicative of microcardia (**Fig 7I,J; VideoS1**). Of note, this double knockdown phenotype is worse than *NP>Phb1 RNAi* alone (**Fig 7G,I,J**), suggesting a synergistic effect. *NP>Phb2 RNAi* alone also produces severe microcardia (**Fig 7H,J**). To assess heart function, we measured cardiac output, which takes into account the amount of blood (lymph) pumped per unit time (stroke volume), the heart chamber size, and the heart rate (see methods). Hypertrophic *NP>COX7A RNAi* hearts exhibit abnormally elevated cardiac output (**Fig 7K**) whereas the microcardia observed in *NP>Phb1 RNAi;COX7A RNAi* double knockdown hearts correlates with reduced output (**Fig 7K**), similar to *NP>Phb1 RNAi* or *NP>Phb2 RNAi* single knockdown (**Fig 7K**). These findings highlight the requirement of mitochondrial integrity for COX7A-mediated repression of cardiac hyperploidy. In summary, our work here reveals how tissue-specific rewiring of mitochondrial regulation controls optimal cell and organ growth in the important context of heart ploidy control (**Fig 7L,M**).

## DISCUSSION

Optimal cell and organ size ensures proper function. To achieve optimal cell size during organ growth, cytoplasmic and nuclear growth processes must be coordinated. Here, focusing on nuclear-based ploidy control, we reveal a unique growth repressor role for COX7A among all annotated ETC genes. This growth repression is conserved in cardiac tissue between flies and humans. COX7A switches from functioning as growth promoter to a suppressor in cardiac tissue due to both a sensitivity of cardiomyocytes to upregulate cytoplasmic mitochondrial biogenesis when mitochondrial regulation is altered (e.g. COX7A or ETC gene knockdown), and a heart-specific lack of COX7A control over mitochondrial integrity (independent of core ETC function). As a result, COX7A knockdown drives over-production of intact mitochondria (i.e mitochondria with preserved mitochondrial integrity). Due to the positive feedback between mitochondrial production and endoreplication that we highlight here, COX7A cardiomyocytes exhibit nuclear hyperploidy and cellular hypertrophy. These changes alter cardiac output, underscoring a functional impact. Of note, mitochondrial dysregulation (mitochondrial myopathy) can lead to cardiac hypertrophy (Kirichenko et al., 2025; Ranjbarvaziri et al., 2021). Our study highlights a mechanism by which the direction of cytoplasmic and nuclear growth (promotion or repression) can be switched through altering the mitochondrial network. Further, we highlight the importance of tissue context in study of cellular and organ growth control.

### Tissue specific rewiring of the ETC facilitates polyploidization

Our study reveals new regulation of polyploidy through mitochondria by the COX7A protein family. We identify the mitochondrially localized *Drosophila* COX7A isoform (IsoA) as promoting polyploidization in the salivary gland while repressing it in cardiomyocytes. Similarly, we find that the human ortholog of *Drosophila* COX7A-IsoA is COX7A1, a heart-specific isoform (Huttemann et al., 2012). COX7A1 is a marker of cardiomyocyte maturation during embryonic development (West et al., 2018). By contrast, the predominantly nuclear localized *Drosophila* COX7A-IsoB is orthologous to COX7A2L, which we show functions as a promoter of ploidy in both tissues. COX7A2L and COX7A1 have been shown to organize two distinct mitochondrial respiratory chains in certain cancer cells depending upon the activation state of pyruvate hydrogenase complex (Fernández-Vizarra et al., 2022), highlighting the flexibility of the COX7A family to distinct tissue and metabolic states. Here, we find that simultaneous knockdown of both nuclear and mitochondrial isoforms of *COX7A* in *Drosophila* results in cardiac nuclear hyperploidy, cellular hypertrophy, and abnormally increased cardiac output. These findings provide likely new insight into findings of cardiac hypertrophy in *Cox7a1* null mice, which were generated in the *C57BL/6J* background that also harbors a mutation in *Cox7a2l* (García-Poyatos et al., 2024; Huttemann et al., 2012). We note that COX7A functions as a growth repressor in response to Notch overactivation in the eye imaginal disc (Sorge et al., 2020). However, it can also act independently as a growth promoter in the same tissue (Sorge et al., 2020), suggesting that COX7A activity may be modulated in different contexts to either promote or suppress growth.

No other tested ETC gene functions to repress cardiac ploidy-only COX7A. Mutations associated with electron transport chain (ETC) genes cause leigh’s syndrome, a condition associated with both dilated and hypertrophic cardiomyopathy (Baertling et al., 2017; Popoiu et al., 2023). In our study, suppression of several nuclear-encoded ETC genes, as well as genes involved in maintaining mitochondrial integrity (Phb1/Phb2), in the *Drosophila* larval heart leads to reduced mitochondrial integrity, smaller heart chamber lumen size, reduced cardiac output and decreased cardiomyocyte ploidy. These findings suggest that the interplay between mitochondrial integrity and endoreplication to regulate ploidy is essential for proper organ growth. Together, these results highlight that distinct metabolic cues resulting from altered mitochondrial activity can differentially modulate endoreplication, with COX7A being a critical tuner of cardiac growth through mitochondria. These findings have important implications for cardiac development and disease.

Our study also reveals unique control over mitochondrial biogenesis that occurs in the heart upon disruption of annotated ETC gene function. These findings mirror previous reports of increased cardiac mitochondrial biogenesis (García-Poyatos et al., 2024; Huttemann et al., 2012). Interestingly, we find that suppression of many other annotated electron transport chain (ETC) components, such as COX4 and ND-MNLL, also increases heart mitochondrial density. This indicates that ETC disruption broadly promotes cardiac mitochondrial biogenesis. One possible explanation lies in the mitochondrial unfolded protein response (UPR^mt^), a conserved stress response triggered by ETC dysfunction. In *C. elegans*, ETC impairment activates UPR^mt^ through the nuclear translocation of ATFS-1, the homolog of human ATF5, (Haynes and Hekimi, 2022; Nargund et al., 2012; Wang et al., 2019). ETC disruption in the heart drives a compensatory increase in mitochondrial number via UPR^mt^ activation (Liu et al., 2022). As a further connection between the need for feedback between optimal endoreplication and mitochondrial control, hyperploidy of the accessory gland, a prostate-like polyploid *Drosophila* tissue (Box et al., 2024), causes mitochondrial DNA aberrations (Molano-Fernández et al., 2022). Overall, our findings suggest that the cellular response to mitochondrial dysfunction is differentially regulated across tissues and shaped by organ-specific metabolic demands and stress adaptation programs, and that COX7A is uniquely adept among ETC genes to respond to these changing biological conditions.

### Understanding cardiac disease through the lens of organ-specific polyploidy

Our reverse genetic screen in the *Drosophila* larval heart and salivary gland uncovered both shared and tissue-specific regulators of polyploidy. We previously demonstrated that the *Drosophila* heart undergoes polyploidization during larval development (Chakraborty et al., 2023), similar to the salivary glands (Zielke et al., 2013). However, the timing, progression, and the extent of endoreplication differ significantly between these two organs. Notably, most ploidy regulators identified in the salivary gland screen did not overlap with those found in the cardiac screen. This striking lack of overlap suggests that organ-specific mechanisms regulate organ growth to meet distinct developmental and functional demands.

Modulation of ploidy control is a powerful mechanism by which tissues can meet structural and physiological requirements during both development and regeneration. For example, *Drosophila* salivary gland cells undergo approximately 10 genome doublings (Edgar et al., 2014) and produce high amounts of glue proteins essential for puparium formation and attachment (Beno et al., 2024).

Premature polyploidy disrupts *Drosophila* oogenesis (Herriage and Calvi, 2024). In contrast, cardiomyocytes modulate ploidy based on mechanical workload; studies show that polyploidy in failing human hearts can decrease with mechanical unloading via left ventricular assisted device (LVAD) implantation (Rivello et al., 2001; Wohlschlaeger et al., 2010). Polyploid growth has also been implicated in supporting regeneration and restoring organ function after developmental defects or injury (Bailey et al., 2021; Cohen et al., 2018; Derks and Bergmann, 2020; Lin et al., 2020; Wilkinson et al., 2019). These observations suggest that organ-specific nuclear growth programs can also underlie distinct regenerative responses. Therefore, identifying organ-specific ploidy regulators may provide insights into tissue regeneration.

Overall, our screen reveals heart-specific coordination between nuclear and cytoplasmic growth to support optimal heart formation and function.

## METHODS

### Fly Stocks

Flies were raised on standard fly food provided by Archon Scientific. Fly stocks used in this study are: *NP5169-Gal4/Cyo* (Kyoto: 113612) (Chakraborty et al., 2023; Monier et al., 2005), *UAS-mCherry-NLS* (RRID:BDSC_38425), *UAS-GFP-NLS* (RRID:BDSC_4775), *UAS-Mito-GFP* (RRID:BDSC_8442), *UAS-GFP-COX7A-IsoA* (*this study*), *UAS-GFP-COX7A-IsoB* (*this study*)*, UAS-fzr (Gifted by Mary Lilly), UAS-COX7A RNAi* (VDRC: v25550, RRID:BDSC_57572), *UAS-COX4 RNAi* (VDRC: v3923), *UAS-ND-MNLL RNAi* (VDRC: v33892, v29884), *UAS-fzr RNAi* (VDRC: v25550), *UAS-Hpo RNAi* (VDRC:v10416, RRID:BDSC_33614), *UAS-Phb1 RNAi* (VDRC: v12360), *UAS-Phb2 RNAi* (VDRC: v110070), VDRC control (v60000) and BDSC control (RRID:BDSC_36304). All other RNAi lines used in the screen are mentioned corresponding to the genes in **TablesS1 and S2**.

Crosses were raised at 25°C, except when expressing transgenes (29°C). For all BrdU feeding experiments, staged larvae were fed 0.5 mg/mL BrdU (Sigma) in standard fly food for 24 hours at 29°C (pulse-chase). After BrdU feeding, larvae were transferred to standard fly food and dissected at the WL3 stage. For double RNAi experiments PCR was used to determine the presence of RNAi transgene insertions in *UAS-Phb1 RNAi;UAS-COX7A RNAi* flies. All PCR primers used in this study provided in **TableS3**.

To assess the validity of gene knockdowns, mRNA was extracted from heart and salivary gland using the mirVana™ miRNA Isolation Kit (Invitrogen, #AM1561) and converted to cDNA with the iScript cDNA Synthesis Kit (Bio-Rad, Hercules, CA, #170–8891). Quantitative real-time PCR (qRT-PCR) was performed using the Luna Universal qPCR Master Mix (NEB, Ipswich, MA, #M3003) according to the manufacturer’s protocol. cDNA amplification and FAM/SYBR Green fluorescence detection were carried out on a CFX384 Touch Real-Time PCR Detection System (Bio-Rad). Expression levels of target mRNAs were normalized to the housekeeping gene rp49. All qPCR primers used in this study provided in supplementary file (**TableS3**).

To design RNAi-resistant COX7A isoforms against the BDSC COX7A RNAi line RRID:BDSC_57572, four codons (encoding LSVP) within the target region of the RNAi, in the COX domain, were modified without altering the amino acid sequence. GFP tagged RNAi-resistant COX7A isoform sequences were generated through gene synthesis (Twist Biosciences), cloned into a pBID-UASC plasmid (Addgene plasmid #35200) and transformed into NEB 5-alpha competent cells. Plasmids were purified using a ZymoPure II Plasmid Midiprep Kit (Zymo Research). Plasmids were injected into attP40 (2DL) flies (BestGene). Full sequences are provided in **TableS4**.

### *Drosophila* Larval Fixed Imaging

For mitochondrial imaging, tissues were dissected in SC medium (Schneider’s medium (Gibco) supplemented with 10% fetal bovine serum (FBS)) and fixed with 3.7% paraformaldehyde in Schneider’s medium for 20 minutes. For all other fixed tissue imaging, tissues were dissected in 1XPBS and fixed in 3.7% paraformaldehyde with 0.3% Triton X-100 for 20 minutes. Tissues were then rinsed twice with 1× PBS and blocked overnight at 4°C in blocking solution (1× PBS, 0.3% Triton X-100, and 1% normal goat serum). Primary antibody incubation was performed in blocking solution overnight at 4°C. Tissues were then washed in PBST (1XPBS and 0.1% Triton X-100), followed by secondary antibody incubation in blocking solution for 1 hour at room temperature. For EdU experiments, Salivary glands were dissected in SC medium and incubated in fresh SC medium containing 20 µM EdU for 2 hours at room temperature prior to fixation. EdU detection was performed following the manufacturer’s protocol (Click-iT™ EdU Imaging Kit, Thermo Fisher), with tissues incubated in the detection solution for 30 minutes. For BrdU detection, BrdU immunostaining was performed as previously described (Chakraborty et al., 2023). For actin labeling, tissues were incubated overnight in Alexa Fluor® 488 Phalloidin (1:250, A12381, Invitrogen) diluted in blocking solution. Afterward, tissues were washed twice in PBST for 10 minutes each and incubated with Hoechst (1:1000 in PBST) for 30 minutes. Finally, tissues were then washed once with PBST, rinsed with 1× PBS, and mounted using VECTASHIELD (Vector Labs). Primary antibody used was Rat anti-BrdU (1:200, ab6326; Abcam), Mouse anti-ATP5A (1:500, ab14748; Abcam). Secondary antibodies used were goat anti-rat Alexa Fluor 488 (1:500, A-11006; Invitrogen) and goat anti-mouse Alexa Fluor 568 (1:500, A-11004; Invitrogen). Images were acquired using Nikon ECLIPSE Ti2 inverted microscope (Nikon PLAN APO Lambda D 60X/1.42 Oil objective #MRD71670 or Nikon PLAN APO Lambda D 20X/0.8 objective #MRD70270). For Salivary Gland screen, images were acquired using on Leica MZ10 F stereoscope (Leica Plan APO 1.0X objective #10450028). For super-resolution imaging, tissues were mounted using Prolong Glass Antifade Mountant (#P36982, Invitrogen). Images were then acquired on ZEISS Elyra 7 (Pan-Apochromat 63X/1.4 Oil) and image reconstruction was done using a Structured Illumination Microscopy (SIM) algorithm. 3D rendering of mitochondria was performed using Imaris software.

### Ploidy Analysis

Cardiomyocyte and salivary gland ploidy were quantified as previously described (Chakraborty et al., 2023; Clay et al., 2023). Briefly, tissues were dissected in 1X PBS and fixed in 4% formaldehyde. After rinsing, samples were transferred onto a siliconized coverslip, overlaid with a charged slide containing ∼10 µL 1X PBS, and gently squashed. Slides were snap-frozen in liquid nitrogen, and the coverslip was removed with a razor blade. Samples were then placed in 90% ethanol (−20°C), air-dried, and rinsed in 1X PBS. Nuclei were stained with Hoechst (1:1000, 10 min), washed in 1X PBS, and mounted in VECTASHIELD (Vector Labs). For ploidy measurement, testes from adult flies were dissected and processed together with the dissected larval hearts or salivary glands and mounted on the same slide. Z-stack images (1 µm intervals) were acquired using an upright Zeiss AxioImager M.2 microscope (Zeiss Plan-Apochromat 63X/1.4 Oil objective #420782-9900-000). Ploidy analysis was performed in Fiji by normalizing the integrated Hoechst intensity of each nucleus to the median intensity of haploid spermatids imaged under identical settings. Cardiomyocyte and gland nuclei were marked using *NP5169-Gal4>UAS-mCherry-NLS*. For ploidy measurements in the larval heart chamber, cardiomyocytes in A5 and A6 segments were quantified for ploidy analysis. For the heart ploidy screen (**TableS1: “Tab1-Ploidy Screen”**), two animals per genotype were examined, with a total of 32 cardiomyocytes analyzed. For salivary gland ploidy measurements, posterior gland cells were quantified.

### TMRM and 2-NBDG Staining and Analysis

For TMRM (T668; Invitrogen) and 2-NBDG (N13195; Invitrogen) staining, salivary glands and cardiac organ were dissected in SC medium (Schneider’s medium (Gibco) supplemented with 10% fetal bovine serum (FBS)). The mitochondrial membrane potential probe TMRM produces lower fluorescence intensity than TMRM but causes less inhibition of the ETC (Perry et al., 2011). Tissues were incubated with either 1 µM TMRM or 0.3DmM 2-NBDG in fresh SC medium. Incubations were performed at room temperature on a rocker, 1 hour for TMRM and 30 minutes for 2NBDG. Following staining, tissues were rinsed with Schneider’s medium and transferred to a grooved glass slide, which prevents the tissue from being compressed by the coverslip. The grooves were created using a rubber cement (Fixogum rubber cement, MP Biomedicals, #11FIXO0125). For TMRM fluorescence analysis, multiple single Z-section images were acquired from different regions of the heart chamber and posterior salivary gland cells using a Nikon ECLIPSE Ti2 inverted microscope (Nikon PLAN APO Lambda D 60X/1.42 Oil objective #MRD71670). We used Fiji software to analyze the fluorescence intensity of TMRM. Since TMRM staining is mitochondrial and not uniformly distributed, images were first thresholded to create 8-bit binary images. A selection mask was then generated from the thresholded image and used as a region of interest (ROI) to measure TMRM intensity in the original image. TMRM fluorescence intensity for each image was then normalized to the area of the ROI. For 2-NBDG fluorescence analysis, complete Z-stacks of the heart chamber were acquired using a Nikon ECLIPSE Ti2 inverted microscope (Nikon PLAN APO Lambda D 60X/1.42 Oil objective #MRD71670), while salivary gland Z-stacks were acquired using the (Nikon PLAN APO Lambda D 20X/0.8 objective #MRD70270). Sum intensity projections of the Z-stacks were generated in Fiji. A region of interest (ROI) was selected for each tissue using the selection tool, and 2-NBDG fluorescence intensity within each ROI was normalized to its area. To calculate fold changes between control and experimental groups, each imaging session included a control heart or salivary gland tissue mounted on the same slide as the experimental sample.

### OCT Measurements of *Drosophila* Larval Hearts

Cardiac function of *Drosophila* WL3 cardiac organs was measured using Thorlabs’ Ganymede™ OCT Systems (#GAN632). The resolution of the OCT system is <3.0µm. To image live larval hearts, animals were quickly rinsed in distilled water and transferred to an adhesive platform made from transparent tape (3M Heavy Duty Scotch tape) attached to a glass slide. Each larva was placed ventral side down, with the dorsal side exposed for OCT imaging. Videos of the posterior heart were acquired using the ThorImage OCT 5.6.0. Each video was acquired at a speed of 100 frames/sec for 5 seconds duration. Multiple M-Mode transverse and sagittal real-time videos were recorded for the heart and the aorta chamber. OCT analysis was performed as previously described (Chakraborty et al., 2023).

### S2 cell Transfection

*Drosophila* S2 culture and transfection were performed as previously described (Rogers et al., 2002). Briefly, S2 cells were maintained in Schneider’s *Drosophila* medium (Gibco) supplemented with 10% FBS (Gibco) and penicillin/streptomycin. For each experimental group (pBID-UASC-COX7A IsoA-GFP or pBID-UASC-COX7A IsoB-GFP), 1 X10^5^ cells were seeded and co-transfected with a mito-mCherry plasmid and an inducible metallothionein-Gal4 construct using TransIT®-Insect Transfection Reagent (Mirus, #MIR 6105). Forty-eight hours post-transfection, the metallothionein promoter was induced with 100 µM copper sulfate for 16 hr. For imaging, cells were seeded onto acid-washed No. 1.5 coverslips (Corning) pre-coated with 0.5 mg/mL concanavalin A (Sigma-Aldrich) in water and air-dried prior use.

### hiPSC-CM Differentiation and Culture

hiPSCs were differentiated into cardiomyocytes (CMs) as previously described (Jackman et al., 2016; Shadrin et al., 2017). Cells were cultured on standard tissue culture dishes or Aclar® coverslips (Ted Pella #10501) coated with Matrigel® hESC-Qualified Matrix (Corning #354277). For plating, hiPSCs were dissociated with Accutase (Stem Cell Technologies) and seeded at a density of 0.64 × 10^6 cells/cm² in mTeSR+ medium (Stem Cell Technologies) supplemented with 5 µM Y-27632 (ROCK Inhibitor; Tocris #1254) for 24 h, followed by mTeSR+ without Y-27632 for 48 h prior to induction of differentiation. On the day of differentiation, cells were incubated with RPMI-1640 medium supplemented with B27 minus insulin (RB-; Thermo #A1895601), 60 ng/mL activin A (Peprotech #120-14E), 12 µM CHIR99021 (Tocris #4423), and 50 µg/mL L-ascorbic acid 2-phosphate (Sigma #A8960) for 24 hrs. Next, cells were maintained in RB-media containing 50 µg/mL ascorbic acid and 5 µM IWR-1 (Tocris #3532/10) for 2 days. Ascorbic acid was withdrawn on day 4. On day 6, cells were switched to RPMI-1640 with B27 plus insulin (RB+; Thermo #17504044) for 48 hrs. On day 8, metabolic selection was initiated using glucose-free RPMI (Thermo #11879020) supplemented with 4 mM lactate (Sigma #L4263), 0.5 mg/mL recombinant human albumin (Sigma #A6612), and 213 µg/mL L-ascorbic acid 2-phosphate (Sigma #A8960) for 48 hrs (Burridge et al., 2014). Thereafter, CMs were maintained in 3DRB+ medium (RB+ supplemented with 2 mg/mL aminocaproic acid (Sigma #A2504), 50 mg/mL ascorbic acid (Sigma #A8960), 0.45 µM thioglycerol (Sigma #M6145), 1% Pen/Strep (Thermo #15070063), 1% non-essential amino acids (Thermo #11140050), and 1% sodium pyruvate (Thermo #11360070)), as previously described (Shadrin et al., 2017), until use.

### Lentivirus Production

Lentiviral plasmids containing U6 shRNA and hPGK eGFP inserts were ordered from VectorBuilder (Scramble Vector ID #VB010000-0001mty, COX7A1 Vector ID #VB240507-1263zvr and COX7A2L Vector ID #VB240507-1265wxx). Lentiviruses were generated as previously described (Nguyen et al., 2016). HEK293T cells (ATCC) were seeded and cultured to 60–80% confluence prior to transfection. Cells were transfected with either Scramble shRNA, COX7A1 shRNA, COX7A2L shRNA or MHCK7 H2B-mCherry reporter plasmids, along with psPAX2 (packaging plasmid) and pMD2.G (envelope plasmid) at a 2:1:1 mass ratio, using JetPrime transfection reagent (PolyPlus). The culture medium was replaced 12-16 hrs after transfection, and virus-containing supernatants were harvested between 48 and 72 hrs post-transfection. To concentrate viral particles, supernatants were mixed at a 3:1 ratio with PEG solution (40% PEG 8000, 24 g/L NaCl), incubated overnight at 4 °C, and centrifuged at 1500 × g for 45 min. The resulting viral pellets were resuspended and stored at-80 °C until use.

### Ploidy Analysis in hiPSC-CMs

After 5 days post virus dosing, hiPSC-CMs were analyzed for Hoechst intensity by flow cytometry. MHCK7 H2B-mCherry and shRNA eGFP lentiviruses were applied at a MOI of 0.8-1. Cells were trypsinized for 3.5 minutes prior to fixation with 2% PFA at RT for 10 minutes. Cells were subsequently incubated with Hoechst 33342 prior to analysis. Cells were analyzed on a FortessaX20 flow cytometer. Using FlowJo software, cells were gated for GFP (shRNA+) and H2B mCherry (CMs), and Hoechst (DNA content). Percentage (%) polyploidy was calculated by combining all cells with ploidy >2n into a single polyploid population.

### <u>hiPSC</u>-CM immunostaining and TMRM assay

Cells were seeded onto Aclar® Coverslips (Ted Pella #10501) prior to differentiation or shRNA experiments. For fixed cell immunostaining, cells were washed once with 1× PBS and fixed with 4% paraformaldehyde for 10 minutes at room temperature (RT). After fixation, samples were washed twice with 1× PBS and incubated in blocking solution (1× PBS containing 0.1% Triton X-100 and 5% donkey serum) for 40 minutes at RT. Cells were then incubated with Hoechst in blocking solution for 30 minutes at RT, followed by three 10-minute washes with PBST (1× PBS containing 0.1% Triton X-100). Finally, samples were mounted with ProLong Glass Antifade Mountant (Thermo #P36984). For Live cell imaging, cells were stained with TMRM for 1 hour, followed by Hoechst staining for 10 minutes. Cells were then imaged on an Andor Dragonfly 505 unit equipped with Borealis illumination spinning disk confocal microscopy, using a Z-step size of 0.25 µm.

Images were captured with an iXon Life 888 EMCCD camera and a 63x/1.47 TIRF HC PL APO CORR oil objective (Leica 11506319; working distance 0.10 mm).

### Phylogenetic Analysis

Protein domain analysis was performed with the InterPro online tool at EMBL-EBI (Blum et al., 2024) to identify Cyt_c_Oxidase_VIIa domains. Sequences were then separated into N-terminal regions and Cyt_c_Oxidase_VIIa domains (**TableS2-“TargetP-2.0 scores and Sequence”**). Next, the N-terminal regions were used for % Identity Matrix analysis with Clustal 2.1. Since the N-terminal region of *Drosophila* COX7AL2 contained only two nucleotides, it was excluded from the % Identity Matrix analysis (**TableS2-“%Identity Matrix-N-terminal”**). Next, mitochondrial targeting signals in COX7A isoform sequences were predicted using the TargetP-2.0 online platform (Almagro Armenteros et al., 2019). Full-length protein sequences were analyzed, and results were classified into two groups: sequences with high mitochondrial targeting probability (>0.7) and those with low or no signal (<0.7) (**TableS2-“TargetP-2.0 scores and Sequence”**). Since the N-terminal region of *Drosophila* COX7AL shared less than 20% identity with COX7A1, it was excluded from the cluster containing mitochondrial targeting signals. Next, alignments for each cluster (high and low mitochondrial targeting probability) were generated using Clustal Omega (Madeira et al., 2024). Finally, the alignment files were used for phylogenetic analysis with the Simple Phylogeny tool at EMBL-EBI (Madeira et al., 2024).

For cladogram analysis, human Complex IV gene orthologs were manually searched in the NCBI Orthologs database (O’Leary et al., 2024) for the following species: *Macaca mulatta, Mus musculus, Rattus norvegicus, Danio rerio, Xenopus tropicalis, Drosophila melanogaster, Anopheles gambiae, Caenorhabditis elegans, Saccharomyces cerevisiae, Arabidopsis thaliana, and Escherichia coli*. A presence/absence matrix (“Yes/No”) was generated from these searches and subsequently used to construct the cladogram.

### Statistics

Statistical methods of analysis, number of biological replicates, values of n, and P values are detailed in the figure legends. Statistical analysis was performed using GraphPad Prism 10.2.3. Statistical notations used in figures: (P > 0.05, (ns) not significant); (P < 0.05, *); (P < 0.01, **); (P < 0.001, ***); (P ≤ 0.0001, ****).

## ACKNOWLEDGMENTS

We thank Bloomington *Drosophila* Stock Center, Vienna *Drosophila* Resource Center and Dr. Zhao Zhang for providing reagents. Dr. Jessica Sawyer provided manuscript comments. Dr. Lisa Cameron, Dr. Yashen Gao, Benjamin Carlson and the Duke Light Microscopy Core provided imaging assistance.

Project support by author: A.C.-National Heart, Lung, and Blood Institute (NHLBI) grant (K99HL177179-01A1) and an American Heart Association (AHA Award ID 23POST1013432), S.D.-an NIH predoctoral fellowship (F31HL162460), N.B.-a National Heart, Lung, and Blood Institute (NHLBI) grant (R01HL164013, R01HL160654 and U01HL134764), S.R.-a bridge funding provided by Integrative Program for Biological and Genome Sciences and support from Stephen Crews and Mark Peifer., D.F.-a National Institute of General Medical Sciences grant (R01GM118447)

