## Supplemental Figures and Legends for "Organ-specific rewiring of mitochondrial integrity through COX7A dictates cellular ploidy control"

**Supplementary files include**

1. **7 Supplementary Figures**
2. **1 Supplemental Movie**
3. **4 Supplemental Tables**

**SUPPLEMENTARY FIGURES:**

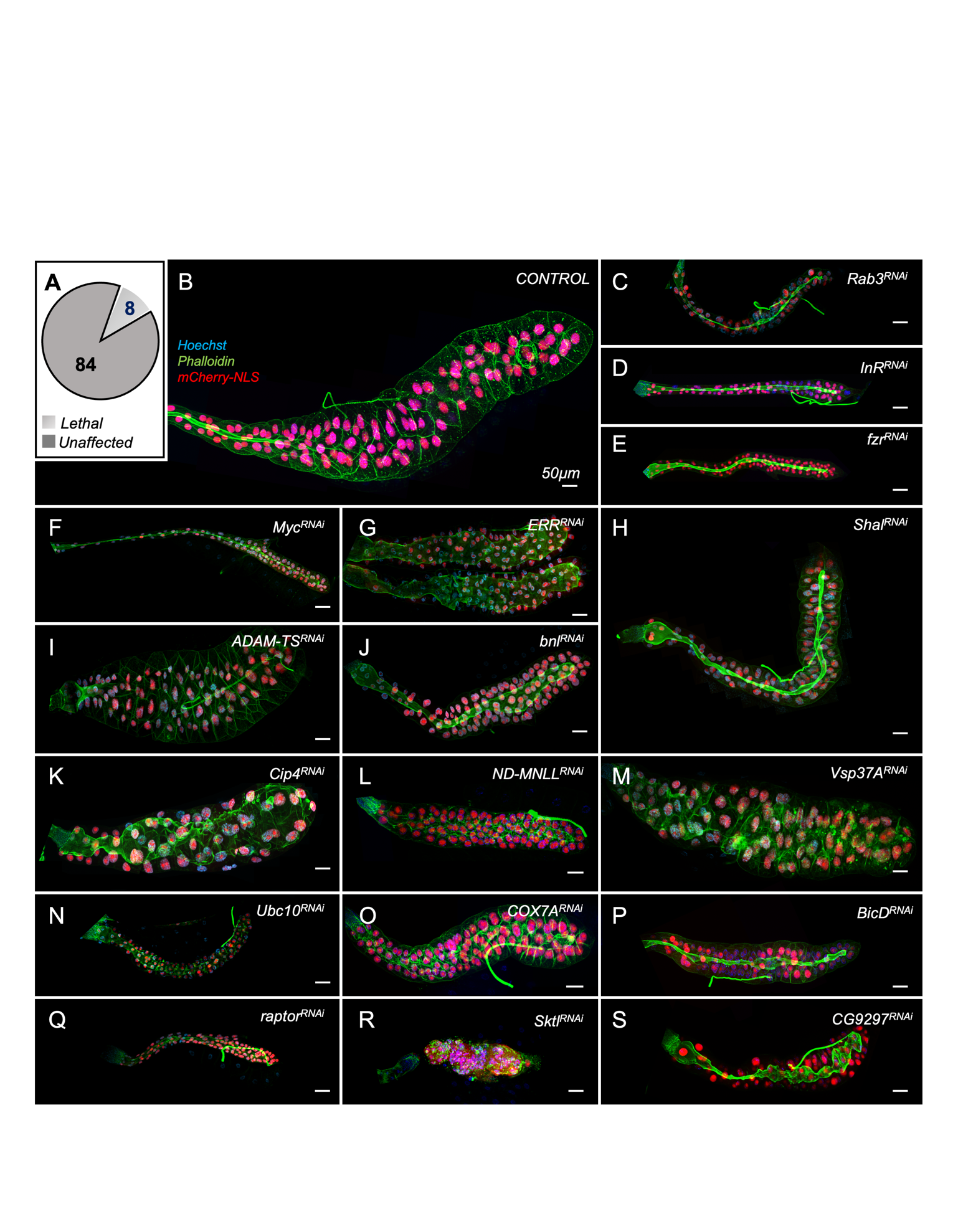

**Figure S1:** **Ploidy regulators of *Drosophila* larval salivary glands**

**(A)** Venn diagram showing the total number of lethal and unaffected genotypes from the ploidy screen. UAS-RNAi lines targeting individual genes in the heart and salivary glands are expressed using *NP5169-Gal4,UAS-mCherry-NLS/CyO* (*NP>*). **(B-S)** Representative salivary gland images of the control (**B**) and the positive regulators of salivary gland ploidy (**C–S**). Gland wall (green, Phalloidin staining), gland nuclei (magenta, *NP>UAS-mCherry-NLS* and Hoechst), and non-gland nuclei (blue, Hoechst).

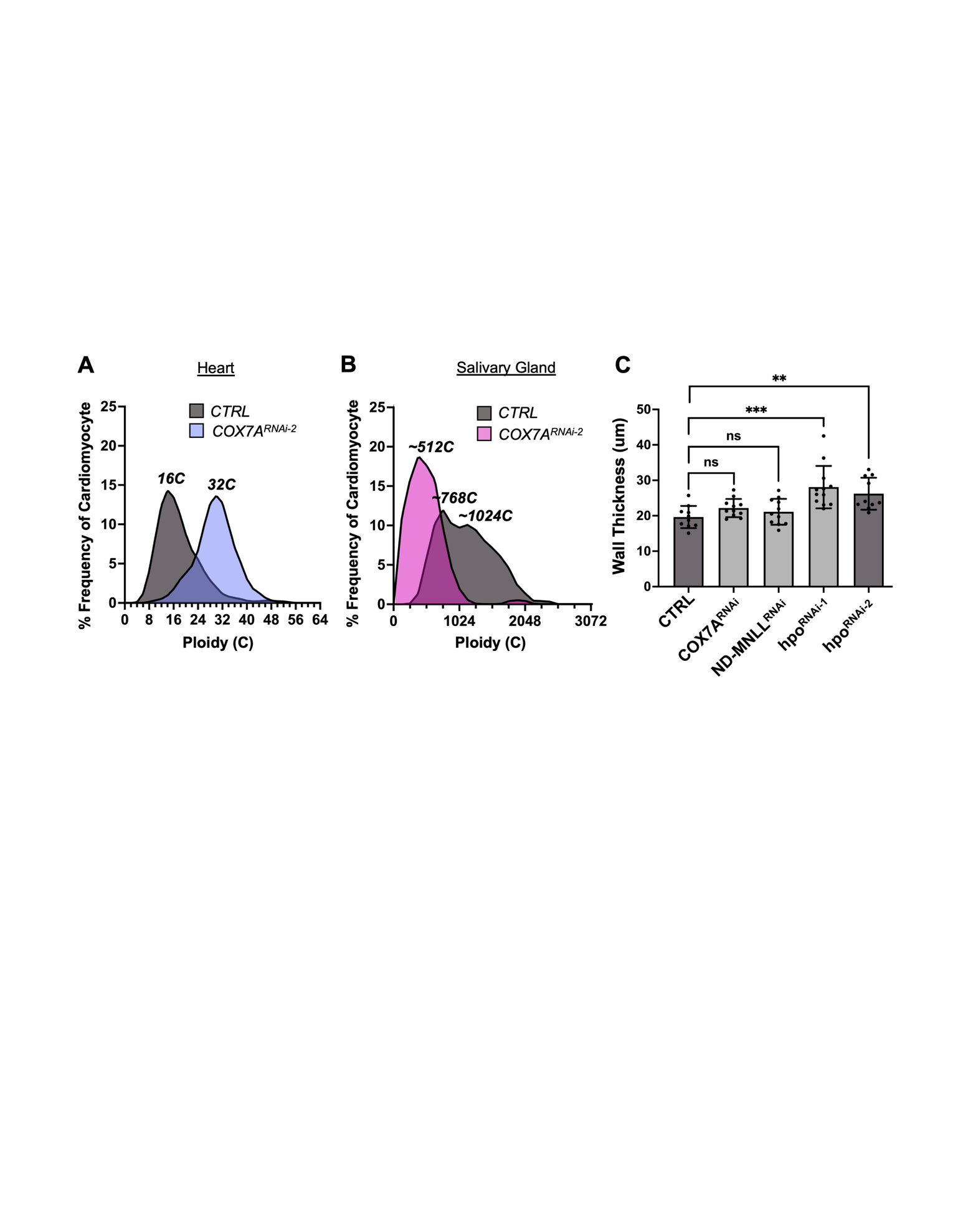

**Figure S2: COX7A regulates nuclear ploidy in tissue-specific manner.**

**(A-B)** Ploidy distribution in control and *NP>COX7A-RNAi* hearts (**A**) and salivary glands (**B**), using a second RNAi targeting the *COX7A* gene. n>5 animals/group. For each animal, the ploidy of 16 cardiomyocytes in the heart chamber (A6–A5) and at least 20 posterior salivary gland cells were analyzed. Each data set includes at least two biological repeats. **(C)** OCT measurements of total wall thickness in the WL3 heart chamber for control (*NP>V60000*), *NP>COX7A-RNAi*, *NP>ND-MNLL-RNAi*, *NP>hpo-RNAi 1* and *NP>hpo-RNAi 2*. Mean±SD; ^ns^P>0.05, **P<0.01, ***P<0.001; Unpaired two-tailed Student’s t-test, n ≥ 10 per group. Each data set includes at least two biological repeats.

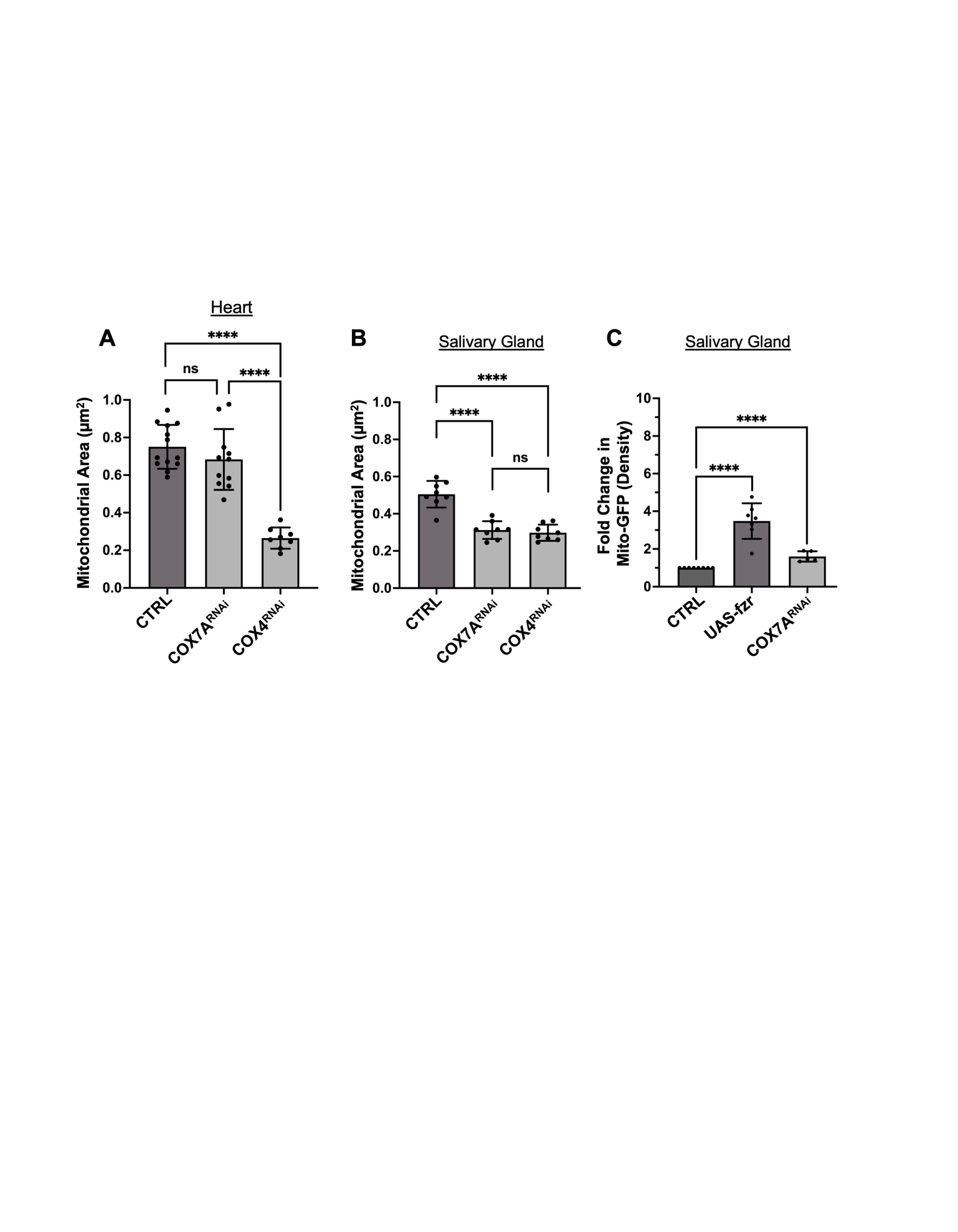

**Figure S3: Tissue-specific mitochondrial regulation of COX7A.**

**(A-B)** Quantification of mitochondrial area in the heart chamber (**A**) and salivary glands (**B**) for the indicated genotypes. Mean±SD; ^ns^P>0.05, ****P<0.0001; Unpaired two-tailed Student’s t-test, n ≥ 5 animals/group. Each data set includes at least two biological repeats. **(C)** Fold change in fluorescence intensity of *NP>UAS-mito-GFP* labeled mitochondria in salivary glands for indicated genotypes. Mean±SD; ^ns^P>0.05, ****P <0.0001; Unpaired two-tailed Student’s t-test. n ≥ 5 animals per group. Each data set includes at least two biological repeats.

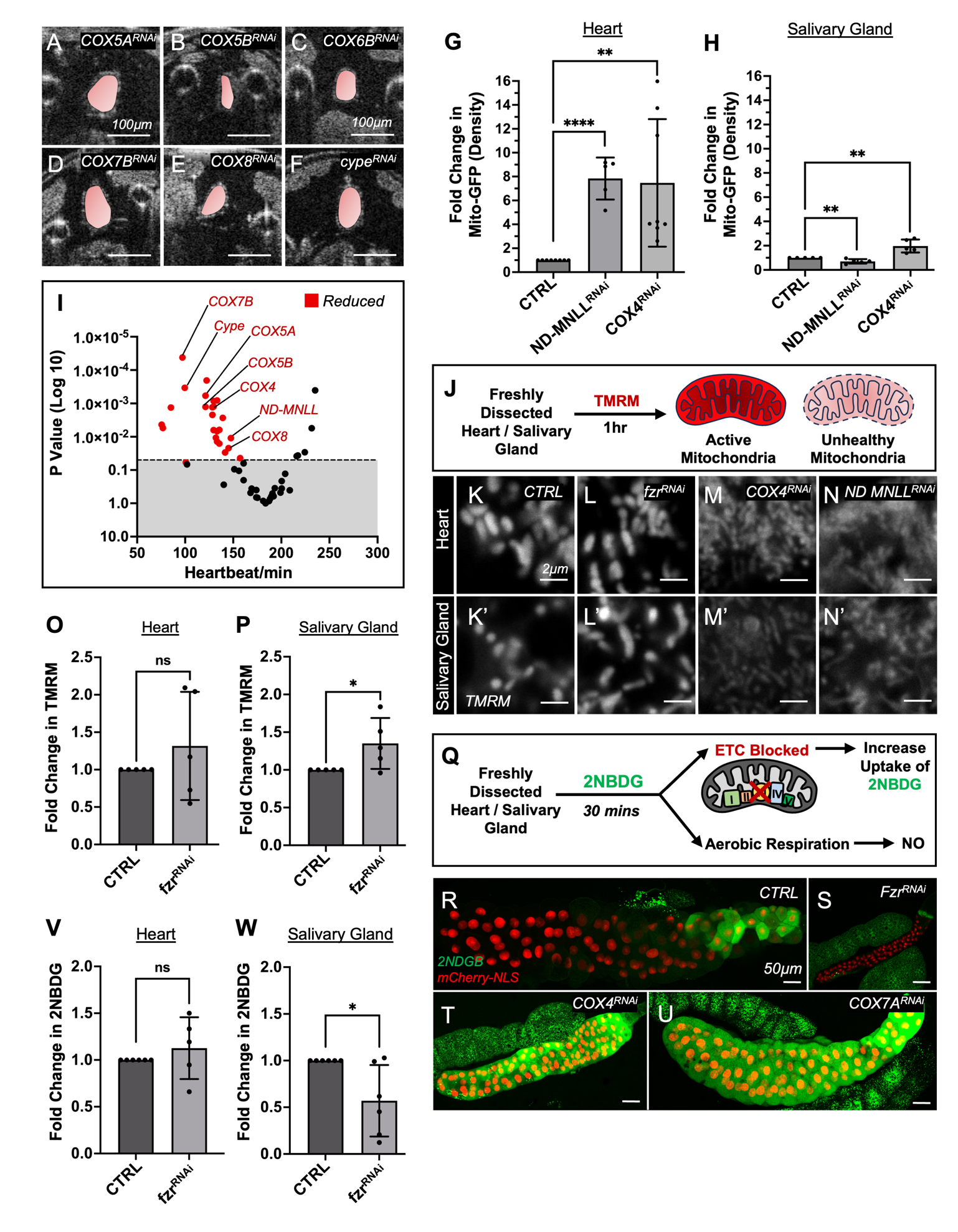

**Figure S4: Impact of ETC and Fzr on mitochondrial integrity and glycolysis.**

**(A–F)** Representative transverse two-dimensional OCT images of WL3 heart chambers showing EDA (pseudo-colored) for the indicated ETC Complex IV gene knockdowns. **(G-H)** Fold change in fluorescence intensity of *NP>UAS-mito-GFP* labeled mitochondria in heart chamber (**G**) and salivary glands (**H**) for the indicated genotypes. Mean±SD; **P<0.01, ****P <0.0001; Unpaired two-tailed Student’s t-test. n ≥ 5 animals per group. Each data set includes at least two biological repeats. **(I)** Volcano plot showing OCT measurements of heartbeat per minute in WL3 heart chambers for 55 electron transport chain (ETC) gene knockdowns. Red dots indicate gene knockdowns that significantly decrease heartbeat (p < 0.05). Mean ± SD; Unpaired two-tailed Student’s t-test. n ≥ 5 animals per group. **(J)** Schematic showing the TMRM-based assay for detecting functional mitochondria in live tissue. **(K-N’)** Representative images of TMRM-stained mitochondria in the heart chamber (**K-N**) and salivary glands (**K’-N’**) for the indicated genotypes. **(O-P)** Fold change in TMRM intensity in control (*NP>V60000*) and *NP>fzr-RNAi* heart chamber (**O**) and salivary gland (**P**). Mean±SD; *P<0.05, ^ns^P>0.05; Unpaired two-tailed Student’s t-test, n>=5 animals/group. Each data set includes at least two biological repeats. **(Q)** Schematic showing the 2NBDG-based assay to detect glucose uptake in live tissue. **(R-U)** Representative images of 2NBDG-labeled (green) salivary glands for the indicated genotypes. **(V-W)** Fold change in 2NBDG intensity in control (*NP>V60000*) and *NP>Fzr-RNAi* heart chamber (V) and salivary gland (W). Mean±SD; *P<0.05, ^ns^P>0.05; Unpaired two-tailed Student’s t-test, n ≥ 5 animals per group. Each data set includes at least two biological repeats.

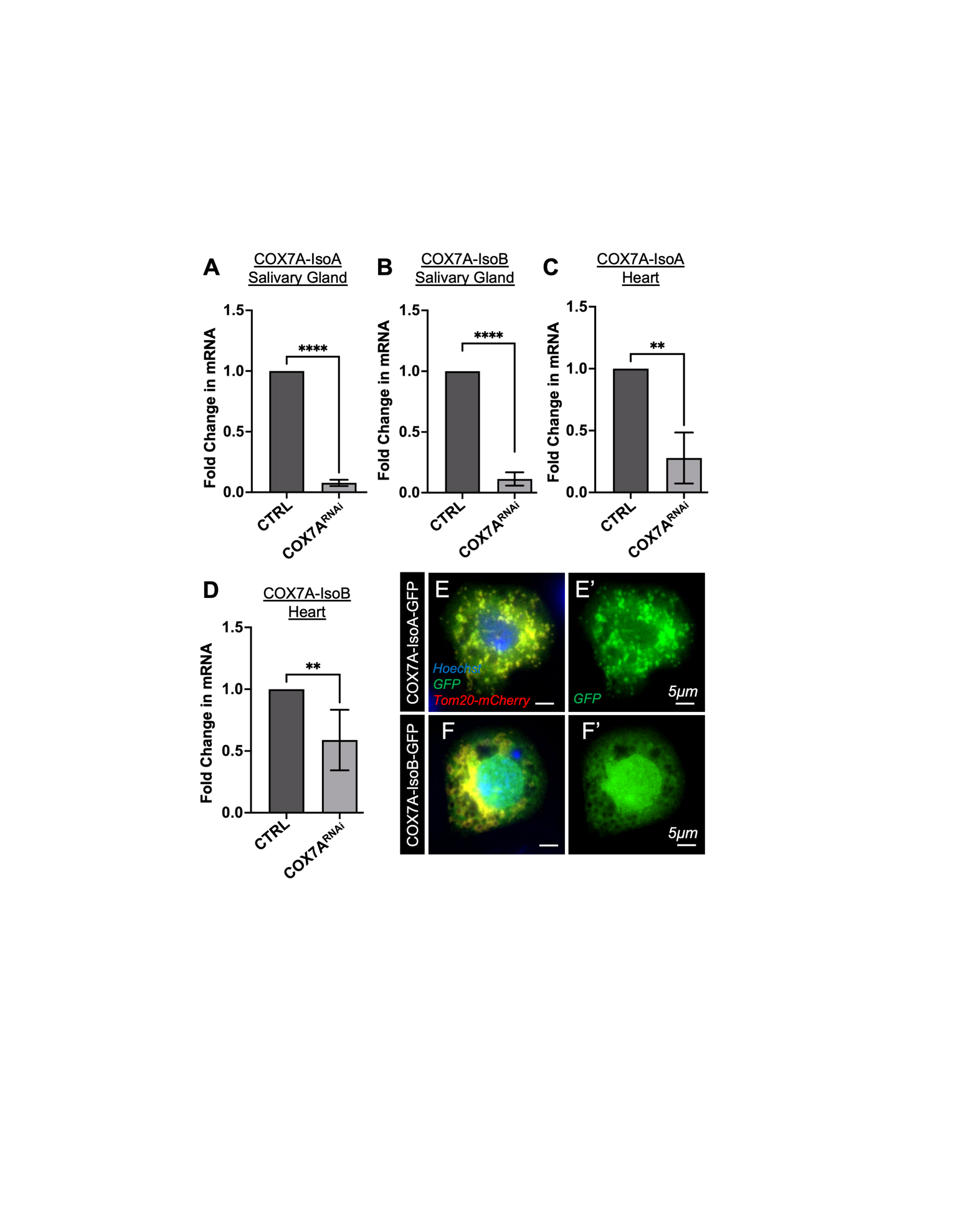

**Figure S5: COX7A RNAi targets both isoforms in heart and salivary glands.**

**(A-B)** Fold change in COX7A Isoform A (**A**) and Isoform B (**B**) mRNA expression in salivary glands. Mean±SD; **P<0.01, ****P<0.0001; Unpaired two-tailed Student’s t-test. All experiments groups have atleast two biological repeats. **(C-D)** Fold change in COX7A isoform A (**C**) and isoform B (**D**) mRNA expression in the cardiac organ. Mean±SD; **P<0.01; Unpaired two-tailed Student’s t-test. All experiments groups have at least two biological repeats. Each data set includes at least two biological repeats. n>10 **(E-F’)** Representative images of S2 cells overexpressing GFP tagged COX7A Isoform A (E-E’; green) and GFP tagged COX7A Isoform B (**F-F’**; green). Mitochondria are labeled with Tom20-mCherry (red; **E, F**).

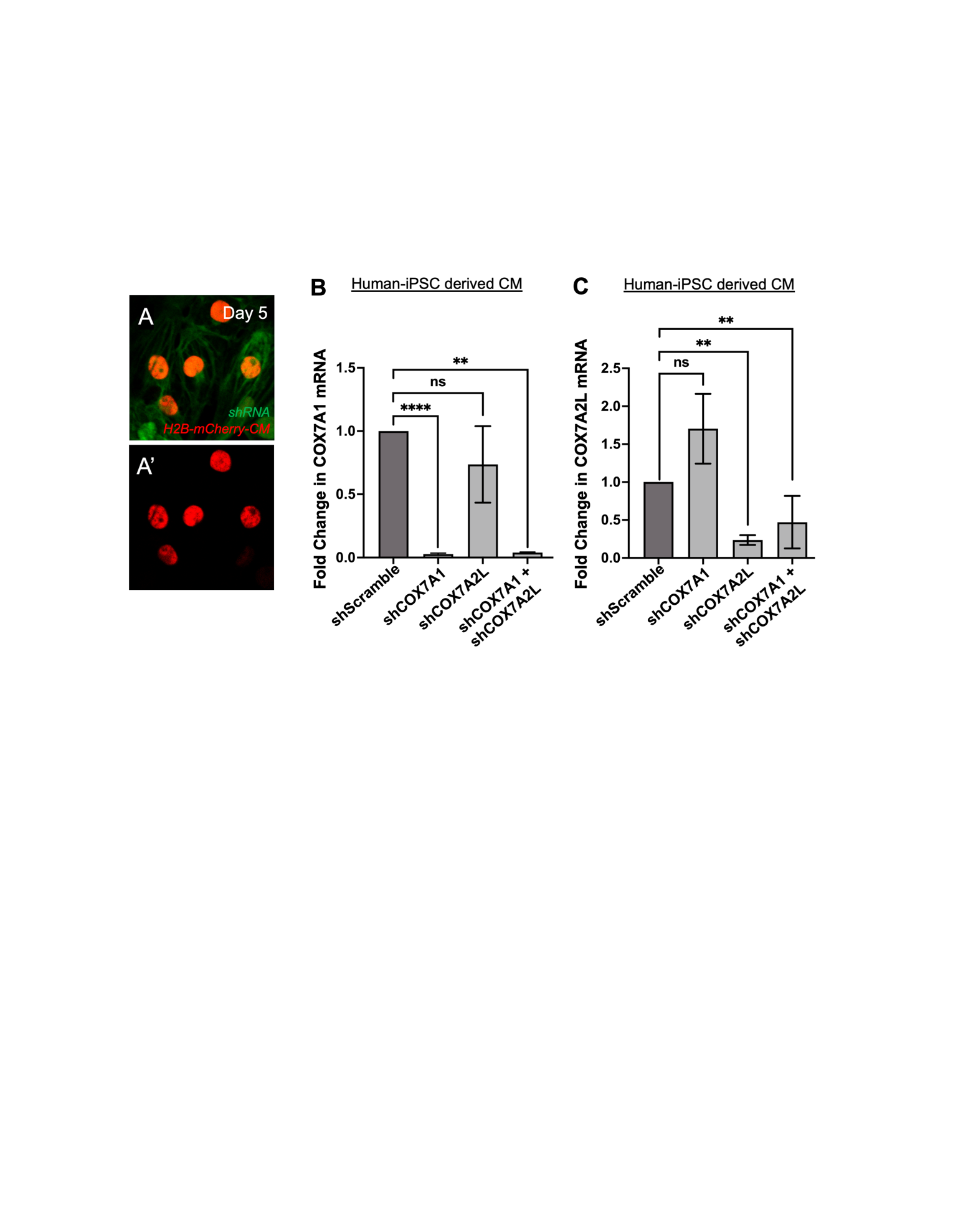

**Figure S6: shRNA mediated knockdown of COX7A1 and COX7A2L in human iPSC-CMs.**

**(A)** Representative images of iPSC-CMs following shRNA-mediated knockdown, labeled with Histone 2B-mCherry (red) and GFP to mark shRNA-expressing cells (green). **(B-C)** Fold change in mRNA expression of human COX7A1 (**B**) and COX7A2L (**C**) in iPSC-CMs following shRNA-mediated knockdown of the indicated genes. Mean±SD; ^ns^P>0.05, ** P < 0.01, ****P <0.0001; Unpaired two-tailed Student’s t-test. Each data set includes at least two biological repeats. 5x10^5^ cells were seeded for each group.

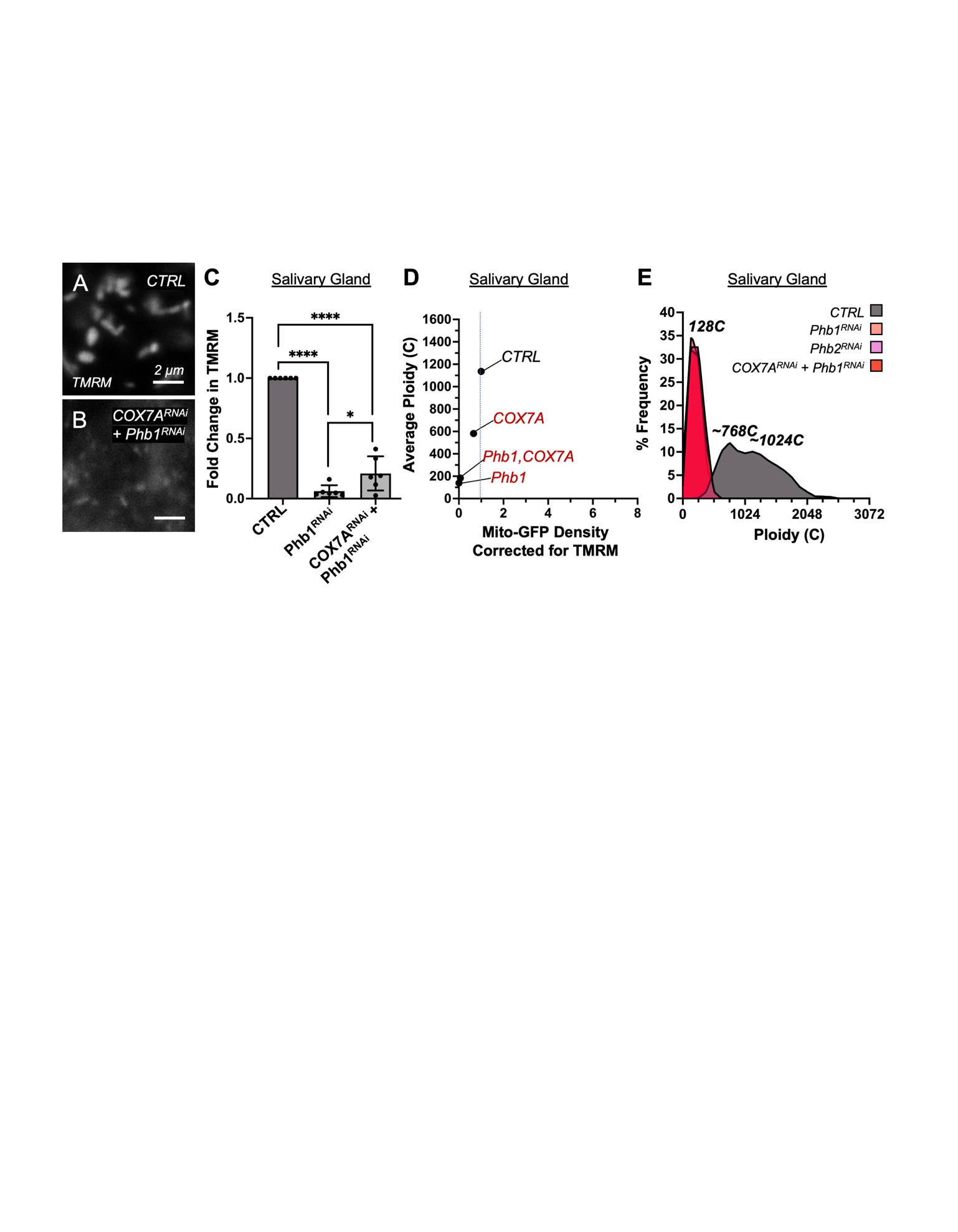

**Figure S7: COX7A-mediated suppression of the cardiac endocycle requires mitochondrial function.**

**(A-B)** Representative images of TMRM-stained mitochondria in the salivary glands of control (**A**; CTRL; *NP>V60000*) and *NP>Phb1-RNAi;COX7A-RNAi* (**B**) animals. **(C)** Fold change in TMRM intensity in the salivary gland for the indicated genotypes. Mean±SD; ****P < 0.0001; Unpaired two-tailed Student’s t-test. n ≥ 5 animals per group. **(D)** Scatter plots showing total mitochondrial GFP fluorescence density (intensity/area) for functional mitochondria versus average ploidy for the indicated genotypes in salivary glands. n ≥ 5 animals per group. Each data set includes at least two biological repeats. **(E)** Ploidy distribution in the salivary glands of indicated genotypes. n>5 animals/group. For each animal, at least 20 posterior salivary gland cells were analyzed. Each data set includes at least two biological repeats.

**Supplemental Table 1: Ploidy and OCT based Screens.** The sheet “Tab1-Ploidy Screen” includes all target genes from our ploidy-based screen of human ventricular upregulated genes (related to Figure 1C), along with their corresponding RNAi reagents, ploidy scores, and gland phenotypes. The sheet “Tab2-Ploidy ScreenHits-OCT based” includes selected hits from the ploidy screen that were further analyzed for chamber lumen size (End Diastolic Area) using OCT. The sheet “Tab3-ETC Screen-OCT based” lists 55 ETC gene knockdowns tested for chamber lumen size by OCT, along with their corresponding RNAi reagents and salivary gland phenotypes. The sheet “Tab4-Heartbeat-OCT based” contains OCT-based heartbeat measurements for the same 55 ETC gene knockdowns.

**Supplemental Table 2: Phylogenetic analysis of COX7A orthologs.** The sheet “TargetP-2.0 scores and sequences” includes the TargetP-2.0 scores (see Methods) for the COX7A orthologs. It also summarizes the N-terminal and COX domain sequences of all COX7A orthologs used for phylogenetic analysis. The sheet “%Identity Matrix–N terminal” includes the % identity matrix calculated using Clustal Omega, based only on the N-terminal sequences.

**Supplemental Table: 3:** qPCR primers

**Supplemental table 4:** Gene sequences of the RNAi resistant *COX7A-IsoA-GFP* and *COX7A-IsoB-GFP*.

**Supplemental Movie 1: Mitochondrial Integrity is required for COX7A mediated repression of cardiac ploidy.** Representative transverse OCT videos of heart chambers for the indicated genotypes: *NP>V60000* (CTRL), *NP>COX7A RNAi*, *NP>Phb1 RNAi* and *NP>Phb1 RNAi;COX7A RNAi*. Related to figure **7F-K**.
